# Spontaneous reinstatement of episodic memories in the developing human brain

**DOI:** 10.1101/2025.09.15.676365

**Authors:** Tristan S. Yates, Bridget L. Callaghan, Jennifer A. Silvers, Michelle VanTieghem, Tricia Choy, Kaitlin O’Sullivan, Lila Davachi, Nim Tottenham

## Abstract

Starting early in development, the hippocampus can support the formation of episodic memories, and yet how the developing brain stabilizes these memories to support their long-term accessibility has not been established. This study (N = 49; age_mean_ = 11.68 years) examined how encoded episodic memories are stabilized in development. Using fMRI and multivariate analyses, we tracked neural reinstatement of newly learned item-location-context associations in youth. During encoding, hippocampus and visual cortex activity predicted later memory success. Crucially, during post-encoding rest, spontaneous pattern reinstatement in the angular gyrus and the pregenual medial prefrontal cortex (mPFC) also predicted memory, with age-related changes in the mPFC and superior frontal gyrus. These findings suggest that episodic memory stabilization during childhood and adolescence is broadly supported by developmental change within regions associated with mature function, advancing our understanding of how different memory stages contribute to how we learn and remember across the lifespan.

## Introduction

Episodic memories (i.e., memories of individual past events and their contextual details, such as what happened, where it happened, and when it happened) form not only the basis of one’s acquired knowledge, but also one’s personal identity (Tulving, 2002). In contrast to other forms of memory, episodic memories are characterized as memories for specific events in which multiple elements of a complex event are recalled. Although infants exhibit rudimentary forms of episodic memory (Bauer et al., 2010; Pascalis et al., 1998; Rovee-Collier et al., 1980), the ability to more richly recall episodes from one’s life continues to develop into childhood and adolescence (Donato et al., 2021; Bevandić et al., 2024). Developmental improvements in episodic memory can result from key changes that unfold at different stages of the memory process, including initial encoding, maintenance/storage through memory consolidation processes, and later retrieval. At present, much of the focus on the neural mechanisms of episodic memory during development has been on the initial memory encoding or later memory retrieval. Here, we build on this existing work and additionally ask how the developing brain stabilizes and maintains episodic memories during the post-encoding consolidation period.

A critical part of a memory’s lifecycle is how its representation is stabilized and maintained in the time between encoding and retrieval. This post-encoding period for memory consolidation is a key aspect of systems consolidation theory (Squire, 1992; Alvarez & Squire, 1994) and multiple trace theory (Nadel & Moscovitch, 1997; Nadel et al., 2000), which both propose that the hippocampus ‘teaches’ the cortex encoded memories for long term storage, but differ in the extent to which the hippocampus is involved in long-term storage (Takehara-Nishiuchi, 2021). One mechanism by which consolidation occurs is through communication between the hippocampus and cortex (Tambini et al., 2010; van Kesteren, Fernandez, et al., 2010; Murty et al., 2017; Tompary & Davachi, 2022; Tambini & D’Esposito, 2020; Schlichting & Preston, 2014) whereby learned information spontaneously reinstates and strengthens memory representations (Wilson & McNaughton, 1994; Peigneux et al., 2004; Frankland & Bontempi, 2005; Squire et al., 2015; Yu et al., 2024). Although consolidation has been shown to occur over long time windows and during offline sleep (Diekelmann & Born, 2010; Maquet, 2001; Stickgold, 2005), there has been increased interest in examining memory consolidation shortly after encoding in the awake state, as this could provide insights into the initiation of memory stabilization processes (Tambini et al., 2010; Tambini & Davachi, 2019; Liu et al., 2022; Joshi et al., 2026).

Inspired by seminal work showing sequential replay of learned spatial patterns in rodents during both sleep and wake states (Skaggs & McNaughton, 1996; Wilson & McNaughton, 1994; Foster & Wilson, 2006; Davidson et al., 2009; Carr et al., 2011), functional neuroimaging in human adults has shown memory reinstatement during awake rest in the hippocampus (Tambini & Davachi, 2013; Gruber et al., 2016; Schapiro et al., 2018; Schuck & Niv, 2019; Yu et al., 2024), visual cortical regions (Staresina et al., 2013; Deuker et al., 2013; Schlichting & Preston, 2014; de Voogd et al., 2016; Tambini & D’Esposito, 2020), and the medial prefrontal cortex (mPFC; van Kesteren, Fernandez, et al., 2010; Tompary & Davachi, 2022; Jimenez & Meyer, 2024; Yu et al., 2024). The mPFC in particular plays an important role in consolidating memories into pre-existing knowledge structures for long-term storage through coordination with the hippocampus (Preston & Eichenbaum, 2013; Tompary & Davachi, 2017), with connectivity and activity in the mPFC related to memory reinstatement even shortly after encoding (van Kesteren, Fernández, et al., 2010; Tompary & Davachi, 2017; Jimenez & Meyer, 2024; Tompary & Davachi, 2022, 2024). However, it is unclear whether and how memory reinstatement occurs in the developing brain, since these cortical regions undergo continuous development into adolescence (Gogtay et al., 2004). Moreover, memories formed during childhood are difficult to recall later in life (Hayne & Jack, 2011), suggesting potential changes in how episodic memories are stabilized for long-term storage and access. Thus, spontaneous reinstatement could provide insights into which stage(s) of the memory process continue to develop into childhood and adolescence to predict changes in episodic memory behavior. Some work has shown cortical reactivation of learned information during retrieval or new learning in developmental populations (Schlichting et al., 2022; Schommartz et al., 2025; Varga et al., 2024) but (to our knowledge) none have looked at spontaneous reinstatement at rest and related this to future memory behavior.

In contrast to consolidation processes, extensive research has investigated how the developing brain encodes and retrieves episodic memories. In particular, developmental changes in episodic memory have been tightly linked to the hippocampus (Keresztes et al., 2018; Donato et al., 2021), a region critically involved in episodic memory in adults (Scoville & Milner, 1957) with notably protracted development (Gogtay et al., 2006; Lavenex & Lavenex, 2013). The computational properties of the hippocampus allow it to support several component processes of episodic memory encoding, including relational binding (linking of information from different modalities into a spatiotemporal context; Davachi, 2004, 2006; Ranganath, 2010) and pattern separation (formation of distinct representations of related memories during encoding; Bakker et al., 2008). Accordingly, changes in episodic memory performance over development have been linked to hippocampal structure (DeMaster et al., 2014; Keresztes et al., 2017; Schlichting et al., 2017; Botdorf et al., 2022; Canada et al., 2019) and function, with greater activity during the encoding and retrieval of remembered versus forgotten information (Ghetti et al., 2010; Sastre et al., 2016; Geng et al., 2019; Tang et al., 2021), even in infancy (Yates et al., 2025). The hippocampus shows a capacity for episodic memory encoding in early development, with continuous refinement during childhood and adolescence contributing to increased memory precision, complexity, and flexibility (Ghetti & Fandakova, 2020). Thus, during middle childhood and adolescence, it may be the case that episodic memory improvement is tied more to post-encoding mechanisms such as spontaneous memory reinstatement outside of the hippocampus.

The current study sought to characterize post-encoding reinstatement as a neural mechanism of episodic memory consolidation during development. Youth (age: *M* = 11.68 years, range = 6.65 – 17.61 years, 26 female) participated in a memory encoding task during functional magnetic resonance imaging (fMRI), with rest runs acquired prior to and following encoding (**Figure 1A**). During memory encoding, participants were instructed to learn the association between items (toys or faces), their contexts (scenes), and their specific spatial locations. Some items shared the same scene contexts, meaning that participants needed to form distinct, relationally-bound mnemonic representations during encoding — in other words, they would need to recall these episodic memories *holistically,* a property of episodic memory recall. Encoding trials differed in the number of items paired with the same scene contexts, allowing exploratory examination of the impact of interference on memory performance and neural activity. Importantly, memory encoding scans were flanked by resting state scans to investigate neural reinstatement. After the MRI scan, participants completed a memory test by choosing amongst familiar distractors the item that previously appeared on the scene and its exact location (**Figure 1B**). The behavior of interest was participants’ ability to answer both the item-context and item-location question correctly, in order to tap into holistic episodic memory (Horner et al., 2015). First, we examined the neural mechanisms of episodic memory encoding during this task. We hypothesized that hippocampal activation during encoding would be related to later holistic recall (i.e., correct item-context and item-location recall) (Davachi et al., 2003), given the need to bind relational information and separate related memories. For spontaneous memory reinstatement at rest, prior work in both rodents and humans has implicated the hippocampus (Tambini & Davachi, 2013; Gruber et al., 2016; Schapiro et al., 2018; Schuck & Niv, 2019) and mPFC (Takehara-Nishiuchi & McNaughton, 2008; van Kesteren, Fernández, et al., 2010; Tambini & Davachi, 2019; Jimenez & Meyer, 2024) — however, because the mPFC is a notably later developing region (Casey et al., 2005; Kolk & Rakic, 2022), it is also possible that neural reinstatement may be absent, elsewhere, or not predictive of memory outcomes in the developing brain.

**Figure 1.**
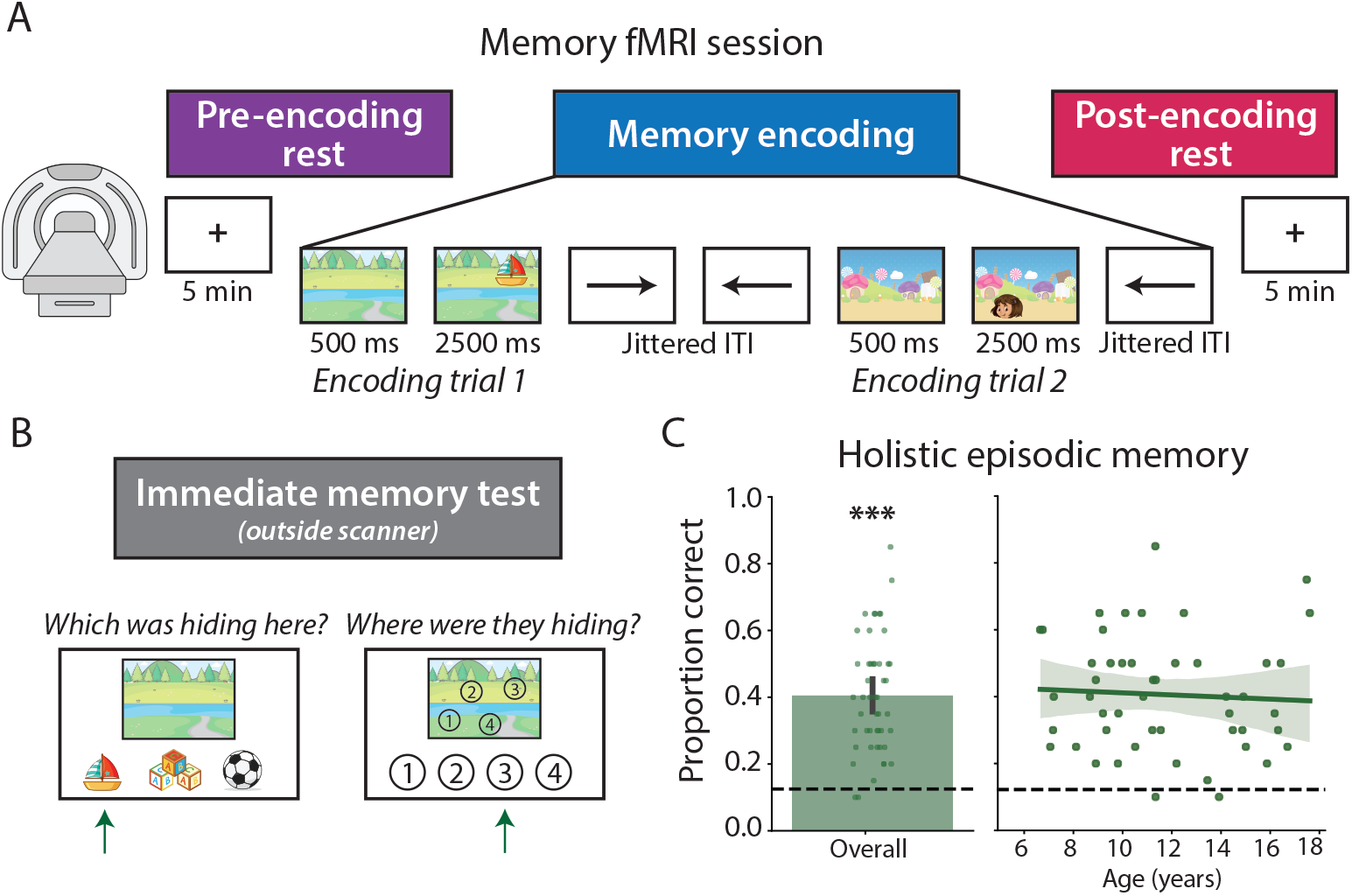
Memory task used to examine episodic memory in a developmental population. (**A**) Participants completed a memory encoding task during fMRI, in which they saw a scene for 500 ms followed by an item (face or object) that appeared in a particular location on that scene for an additional 2500 ms. In between trials, there was a jittered intertrial interval (ITI) in which left or right arrows appeared, and participants were instructed to press any key when they saw an arrow. Prior to and following this encoding run, participants completed 5-minute rest scans in which they stared at a fixation cross. (**B**) Outside of the scanner, participants completed an immediate memory test in which they were asked to choose which item appeared on the scene and where in the scene that item was located. Participants who answered both questions correctly were considered to have ‘holistic episodic memory.’ (**C**) Holistic episodic memory performance overall and across age. Dots represent individual participant data, and error bars and bands reflect the 95% confidence interval from bootstrap resampling. *** *p* < .001.

## Results

### Holistic episodic memory behavior

To measure holistic episodic memory in a developmental sample, memory for item-location-context associations was tested after scanning. Accuracy was defined as correctly recalling both the item-context and item-location associations. Youth showed evidence of holistic episodic memory (*M* = 40.61%, *t*(48) = 11.37, *p* < .001, *d* = 1.64; **Figure 1C**) at performance levels above chance (average chance = 12.5%). Age effects were not observed (*r* = −.056, *p* = .701), consistent with prior work showing that holistic episodic memory emerges by 4 to 6 years of age (Ngo, Horner, et al., 2019; Ngo, Lin, et al., 2019; James et al., 2021). Moreover, participants tended to either remember trials holistically (i.e., the entire item-location-context association) or not at all (**Figure S1**).

### Neural correlates of episodic memory encoding

To examine the neural mechanisms of encoding that underlie holistic episodic memory, a general linear model was performed on preprocessed BOLD activity using separate regressors for trials that were ‘subsequently remembered’ (i.e., the item-location-context association was recalled during the memory test) and trials that were ‘subsequently forgotten’ (i.e., the association was not recalled) (Brewer et al., 1998; Wagner et al., 1998). The contrast of subsequently remembered and forgotten encoding trials was investigated in a whole-brain, voxel-wise manner using a non-parametric randomization approach with multiple comparisons correction (cluster-determining threshold: *z* > 2.41, *p* < .05) and in ROIs used in prior adult research on memory encoding and/or reinstatement as a secondary analysis (**Table S1**).

Youth showed significantly greater activation in the right anterior hippocampus during the encoding of trials that would later be holistically remembered versus forgotten (**Figure 2; Table S2**). This subsequent memory encoding effect was also found in the lateral occipital cortex, bilateral supramarginal gyrus, and left inferior frontal gyrus. No age effects were found in these regions (**Figure S2**); likewise, there were no age effects in the whole brain after multiple comparisons correction. A follow up analysis conducted on ROIs used in prior adult research mirrored these results, with a significant subsequent memory effect in the bilateral hippocampus (**Figure S3**). These results extend prior research showing hippocampal involvement in memory encoding during development (Ofen et al., 2007; Ghetti et al., 2010; Shing et al., 2016) within a complex event requiring holistic episodic memory.

**Figure 2.**
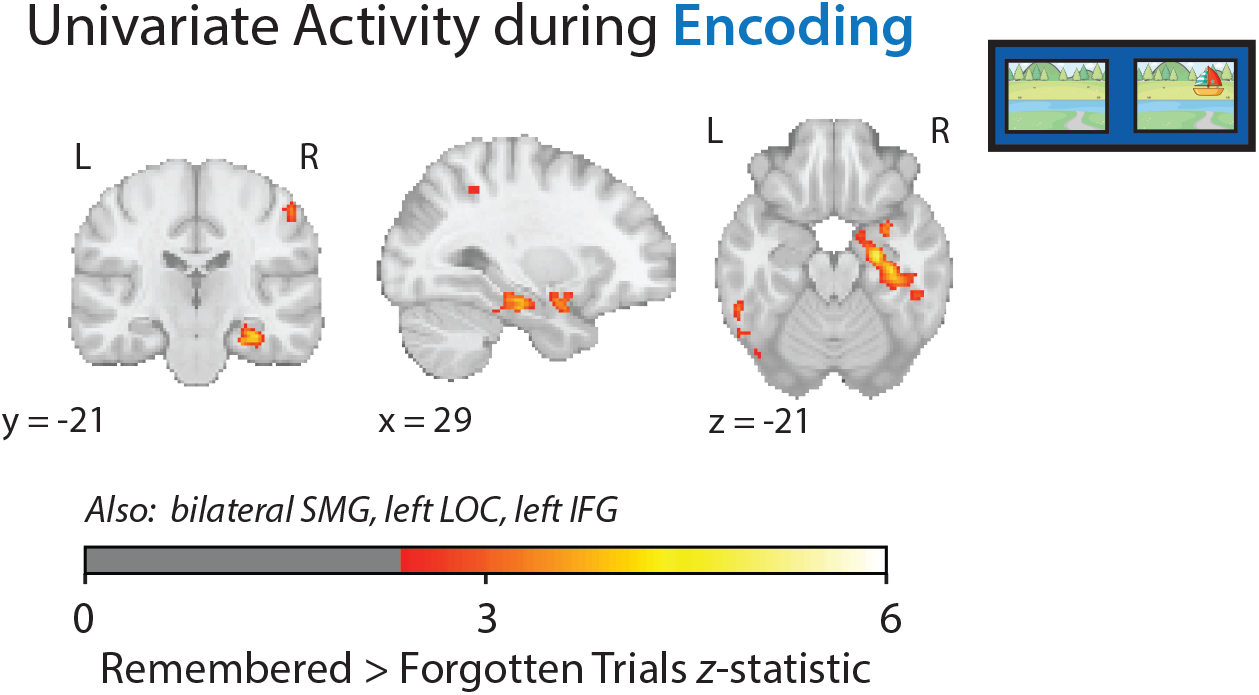
Successful episodic memory encoding effects in youth. Whole-brain univariate activity during memory encoding, showing significant activation of the right anterior hippocampus for holistically remembered versus forgotten encoding trials. Voxels significant following multiple comparisons correction (cluster-defining threshold: *z* > 2.41, *p* < .05) are colored by the average *z*-statistic for the contrast of remembered versus forgotten trials across participants. Significant regions not pictured are listed below the brain plots and are detailed in **Table S2**. SMG = supramarginal gyrus, LOC = lateral occipital cortex, IFG = inferior frontal gyrus.

### Neural correlates of spontaneous memory reinstatement

To ask whether post-encoding memory reinstatement could be detected during development and predict later memory performance, we examined whether trial level brain activity patterns during encoding were reinstated during post-encoding rest and contrasted this with the pre-encoding rest scan, with the logic that higher correlations in post-encoding rest reflect memory reinstatement. We used a multivariate approach to allow us to link activity patterns during encoding with activity patterns throughout the rest period; such patterns at rest cannot be captured in a univariate analysis that requires specified timing. Multivariate analyses were conducted in a whole-brain searchlight by correlating the pattern of activity for each encoding trial with every time point in both the pre-and post-encoding rest scans and then summing these correlations and contrasting between conditions (**Figure 3A**; Staresina et al., 2013; Schapiro et al., 2018; Jimenez & Meyer, 2024).

**Figure 3.**
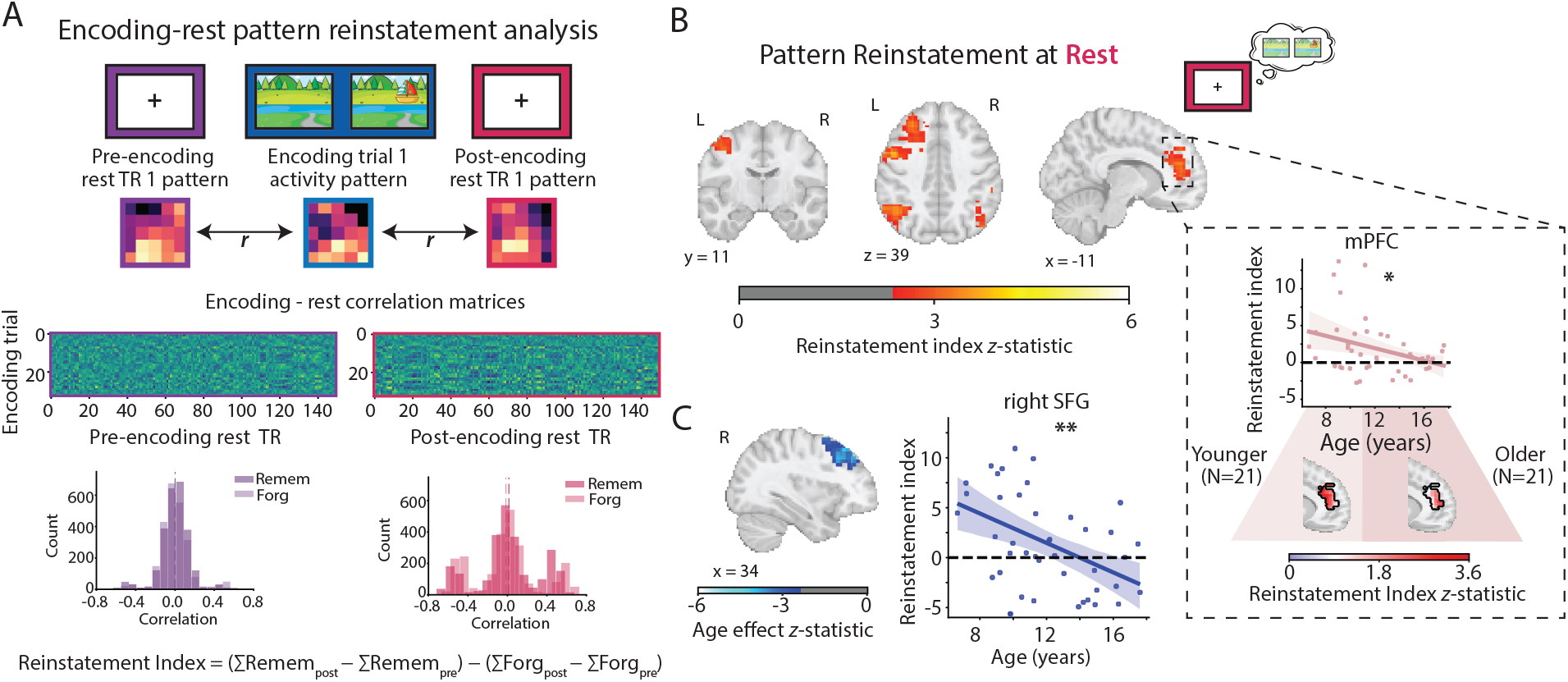
Spontaneous memory reinstatement in youth. (**A**) Analytical approach to examining spontaneous memory reinstatement. Multivoxel patterns of activity during encoding trials were correlated with each TR in the pre-encoding and post-encoding rest runs to construct encoding-rest correlation matrices. These matrices were then summed within each encoding trial type (i.e., holistically remembered or forgotten) and the difference in summed values for remembered versus forgotten trials during post-encoding rest was calculated after accounting for pre-encoding rest. (**B**) Whole-brain multivariate pattern reinstatement at rest, showing significant reinstatement in the medial prefrontal cortex (mPFC), left precentral gyrus, and bilateral angular gyrus. Voxels that were significant following multiple comparisons correction (cluster-defining threshold: *z* > 2.41, *p* < .05) are colored by the average *z*-statistic for the contrast of remembered versus forgotten trials across participants. Inset: Age effects within the mPFC cluster, showing a decrease in the contrast of remembered versus forgotten pattern reinstatement with age. Visualization of the average pattern reinstatement within younger and older children suggest that this may reflect a decrease in the spatial extent of the mPFC contributing to pattern reinstatement over development. (**C**) Whole-brain age effects on pattern reinstatement, showing a decrease in the contrast of remembered versus forgotten pattern reinstatement with age in the right superior frontal gyrus (SFG). Voxels significant following multiple comparisons correction (cluster-defining threshold: *z* > 2.41, *p* < .05) are colored by the average *z*-statistic for the age covariate. ** *p* < .01, * *p* < .05.

During the post-encoding rest scan, youth showed significantly more pattern reinstatement of trials that would later be remembered versus forgotten, controlling for the pre-encoding rest scan, in the bilateral angular gyrus, left precentral gyrus, and pregenual subregion of the mPFC (**Figure 3B; Table S3**). A breakdown of these effects revealed that this was being driven by greater reinstatement of remembered trials during the post-encoding rest scan (**Figure S4**). A follow up analysis conducted on ROIs used in prior adult research mirrored these results, with a significant subsequent memory reinstatement effect in the bilateral angular gyrus (**Figure S5**). Importantly, we validated our approach by showing that these reinstatement effects disappeared when remembered and forgotten encoding trial labels were randomly shuffled within participant (**Figure S6**), and replicated when using the average rather than sum of correlations in the encoding-rest correlation matrices (**Figure S7**).

In contrast to the memory encoding analysis, we found an effect of age on the neural correlates of spontaneous memory reinstatement in youth. Specifically, pattern reinstatement for remembered versus forgotten trials decreased over development in the right superior frontal gyrus (**Figure 3C**). In a median split of children by age, this effect in the right superior frontal gyrus was being driven by younger children, with significant pattern reinstatement of remembered trials in children aged 6 to 11 years (*M* = 3.46; *t*(20) = 3.08, *p* = .006) but not in children aged 11 to 17 years (*M* = −0.55; *t*(20) = −0.75, *p* = .464).

Within the regions that showed an effect of pattern reinstatement in the whole brain, there was an additional age effect in the mPFC, such that reinstatement decreased over development (*r* = −.318, *p* = .04; (**Figure S8A**). Given the mPFC showed significant pattern reinstatement at the group level (unlike the superior frontal gyrus), one possibility is that older children still engaged the mPFC for pattern reinstatement, but in a smaller subregion of the mPFC than younger children. This kind of ‘diffuse to local’ processing has been shown to occur in the prefrontal cortex over development (Casey et al., 2005; Durston et al., 2006). Thus, in an exploratory whole-brain analysis, we visualized spontaneous pattern reinstatement in younger and older children separately; qualitatively, reinstatement within the mPFC was more focal in older children than in younger children (**Figure 3B**), with the number of above-threshold voxels in the mPFC decreasing over development, particularly at higher thresholds (**Figure S8B, C**). These results together suggest changing involvement of the prefrontal cortex in memory reinstatement during childhood and adolescence.

### Modulatory effects of trial type on encoding and reinstatement

Thus far, we have seen that hippocampal encoding and mPFC reinstatement at rest predict children’s memory for item-location-context associations. However, in the current task, items could be located within a scene context that would later be paired with one other item (‘low interference’) or three other items (‘high interference’; **Figure 4A**). This could influence mnemonic processing, since pattern separation may be more difficult when more items share the same scene context. Indeed, although children showed evidence of above-chance holistic episodic memory for both trial types (low interference: *M* = 51.53%, *t*(48) = 8.27, *p* < .001, *d* = 1.19; high interference: *M* = 23.81%, *t*(48) = 6.13, *p* < .001, *d* = 0.89), memory performance was significantly higher when interference was lower (*t*(48) = 6.77, *p* < .001, *d* = 0.98; **Figure 4B**). No age effects were found for trials with lower (*r* = .162, *p* = .268) or higher (*r* = −.122, *p* = .404) interference.

**Figure 4.**
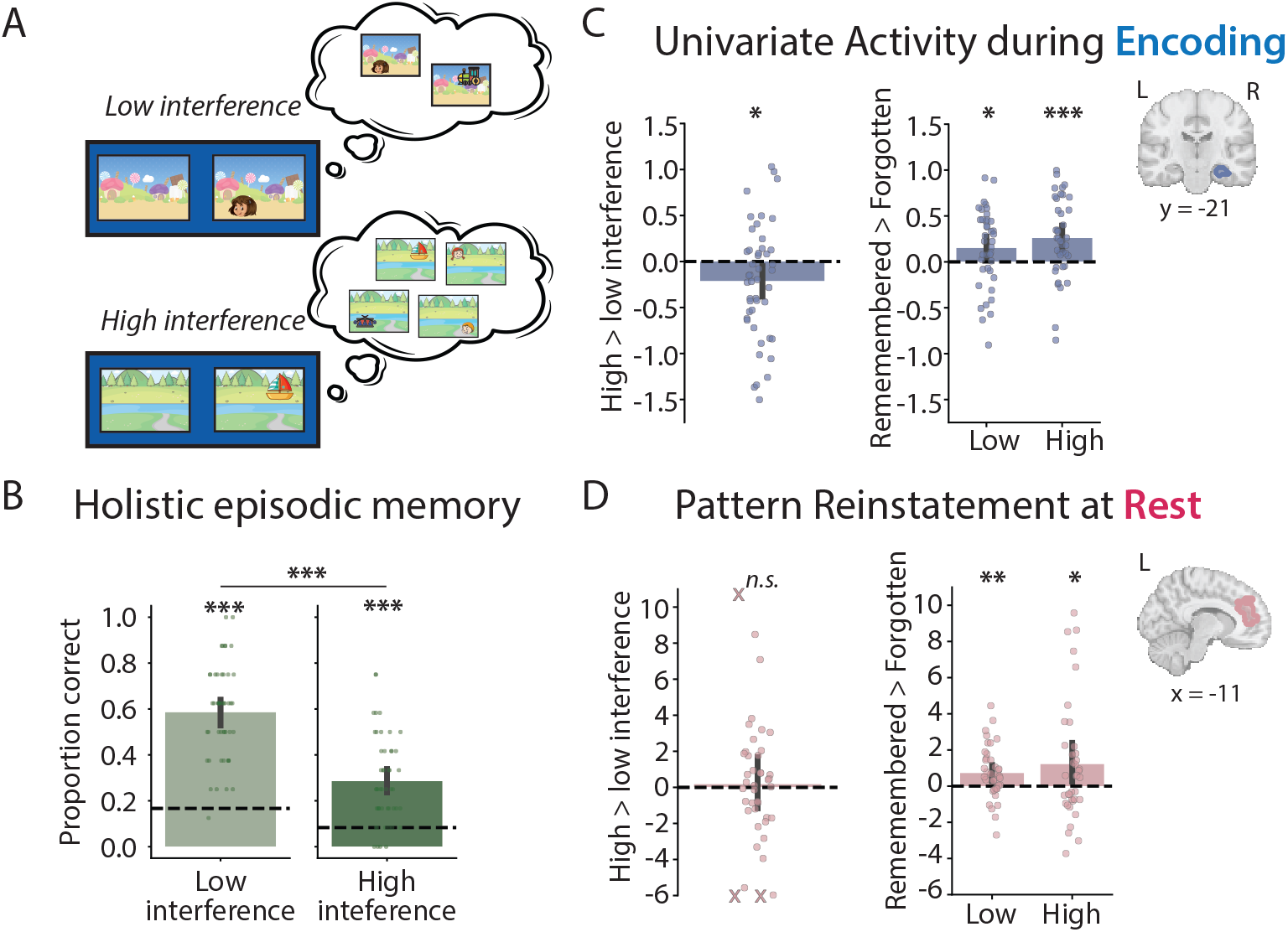
Effects of interference trial type on memory encoding and reinstatement in youth. (**A**) Example low and high interference trials, defined by whether a given encoding trial comprised a scene that would be paired with two or four items. (**B**) Holistic episodic memory behavior by trial type, showing significantly above-chance performance for both trial types and better performance when interference was lower. (**C**) Univariate activity during encoding in the hippocampus, showing less activity during the encoding of high versus low interference trials, but comparable subsequent memory effects. (**D**) Pattern reinstatement in the mPFC, showing no difference in reinstatement of high versus low interference trials and comparable reinstatement related to subsequent memory. Dots represent individual participant data and error bars reflect the 95% confidence interval from bootstrap resampling. Participants with values outside of the plotting range are marked with Xs on the negative or positive edge of the y-axis. *** *p* < .001, ** *p* < .01, * *p* < .05, n.s. = not significant.

We examined the influence of shared scene context on brain activity in two ways. First, we contrasted brain activity during the encoding of low and high interference trials in regions that showed a significant subsequent memory effect in the whole brain. Because participants only ever saw a single item located on a scene at a time (i.e., the structure of visual information was the same regardless of whether a trial was labeled as ‘low’ or ‘high’ interference), this contrast should reflect a memory signal such as interference resolution or pattern separation. There was less activity when youth viewed high interference compared to low interference trials in both the hippocampus (*M* = −0.21, *t*(45) = −2.34, *p* = .024, *d* = −0.35; **Figure 4C**) and lateral occipital cortex (*M* = −0.31, *t*(45) = −3.21, *p* = .002, *d* = −0.48; **Figure S9A**). However, because high interference trials by definition included scenes that repeated more often throughout the task, this effect could also reflect something like repetition suppression for scenes (Kim, 2017). Therefore, in a subset of participants with enough trials per condition, we examined the contrast of holistically remembered versus forgotten trials separately for low and high interference trials. There was significantly greater activity in the hippocampus for remembered versus forgotten trials for trials both low (*M* = 0.15, *t*(39) = 2.27, *p* = .029, *d* = 0.36) and high in interference (*M* = 0.26, *t*(39) = 3.85, *p* < .001, *d* = 0.62), with no difference between trial types in this subsequent memory contrast (*t*(39) = −1.04, *p* = .306, *d* = −0.17). Additionally, there were limited effects of trial type on memory-related encoding activity in other regions that showed an overall subsequent memory effect (**Figure S9B**). These results suggest that the decreased hippocampal activity during high interference trials likely reflects repetition suppression for shared scene contexts.

We performed parallel analyses to examine effects of shared scene context on pattern reinstatement during post-encoding rest. In the mPFC, there was no difference in the reinstatement of low versus high interference trials during post-encoding rest after accounting for pre-encoding rest (*M* = 0.19, *t*(41) = 0.26, *p* = .795, *d* = 0.04; **Figure 4D; Figure S10A**). Moreover, there was significantly greater pattern reinstatement in the post-encoding rest scan for subsequently remembered versus forgotten trials that were both low (*M* = 0.73, *t*(39) = 3.21, *p* = .003, *d* = 0.51) and high in interference (*M* = 1.23, *t*(37) = 2.33, *p* = .025, *d* = 0.38), with no difference between trial types (*t*(35) = −0.40, *p* = .688, *d* = −0.07). There were limited effects of trial type on memory reinstatement in other regions, with the exception of significantly greater reinstatement for remembered versus forgotten high interference trials compared to low interference trials in the left angular gyrus (*t*(35) = 2.09, *p* = .044, *d* = 0.35; **Figure S10B**).

## Discussion

We provide evidence that during childhood and adolescence, the hippocampus encodes episodic memories and the mPFC and angular gyrus spontaneously reinstate encoding patterns during a rest period immediately following encoding. These patterns predict which memories will be subsequently recalled in a holistic manner in the future. The involvement of these regions in encoding and reinstatement mirrors prior work in adults. Additionally, pattern reinstatement of subsequently remembered item-location-context associations decreases over development in the mPFC and right superior frontal gyrus, revealing developmental changes in post-encoding memory processes in the prefrontal cortex.

### Spontaneous episodic memory reinstatement in angular gyrus and prefrontal cortex during development

This study provides developmental evidence of spontaneous reinstatement during an awake consolidation period that predicts later episodic memory recall behavior. Spontaneous reinstatement of learned information during awake rest has been well-documented in the rodent hippocampus (Foster & Wilson, 2006; Diba & Buzsáki, 2007; Karlsson & Frank, 2009; Carr et al., 2011) and more recently shown in adult humans in the hippocampus, mPFC, and cortical regions (Tambini & Davachi, 2013, 2019; Gruber et al., 2016; Schapiro et al., 2018; Schuck & Niv, 2019). Reinstatement may serve different purposes in different regions. For example, reinstatement in cortical regions may strengthen sensory representations; indeed, recent work has shown angular gyrus involvement in item-level reinstatement and memory precision (Hou et al., 2025). In the hippocampus and mPFC, reinstatement may relate to the stabilization of memories for long-term storage, consistent with theories of systems consolidation (Frankland & Bontempi, 2005; Takehara-Nishiuchi & McNaughton, 2008; Preston & Eichenbaum, 2013).

In the current developmental study, we found evidence of spontaneous memory reinstatement during post-encoding rest in regions previously found in adults: the angular gyrus and mPFC. The angular gyrus, part of the posterior medial episodic network (Ritchey & Cooper, 2020), has been implicated in memory retrieval (Rugg & Vilberg, 2013; Wagner et al., 2005), is modulated by vivid remembering (Tibon et al., 2019), and plays a causal role in the retrieval of episodic memory details (Thakral et al., 2017). Thus, one interpretation of our results is that spontaneous memory reinstatement in the angular gyrus relates to holistic episodic memory by enhancing the recall of vivid details. This prompts future work exploring how pattern reinstatement in the angular gyrus relates to vividness to predict the holistic recall of episodic memories. We additionally found reinstatement in the mPFC. This result is striking for several reasons. First, the mPFC is late developing (Kolk & Rakic, 2022), and yet despite this protracted structural development, it appears to be involved in mature function in the current developmental sample. Second, the mPFC is involved in representing abstract generalizations over experiences (i.e., schemas; Tse et al., 2007; van Kesteren, Fernández, et al., 2010; Preston & Eichenbaum, 2013; Robin & Moscovitch, 2017; Tompary & Davachi, 2017; Gilboa & Marlatte, 2017; Baldassano et al., 2018; Tompary & Davachi, 2024). In the current study, reinstatement was related to later memory for specific item-location-context associations that shared contextual information. Participants therefore could have formed an integrated representation of multiple episodes; however, we did not test this behaviorally in this paradigm. Because memory reinstatement can support both episodic memory strengthening and generalization across episodes (Schlichting & Preston, 2014; Tompary & Davachi, 2017; van Kesteren et al., 2010), the mPFC may play an important role in balancing between specific and general memories in development (Forest et al., 2023; Keresztes et al., 2018). Better understanding the functional role(s) of mPFC involvement in awake consolidation during development will require task paradigms that assess memory for both episodic and more generalized, schematic information (Richter et al., 2019; Tompary et al., 2020).

There was an effect of age on spontaneous memory reinstatement in both the mPFC and the right superior frontal gyrus, with younger children showing greater pattern reinstatement of remembered versus forgotten trials during post-encoding rest compared to older children. Importantly, these age effects emerged despite no age-related changes in memory behavior, suggesting that these effects cannot easily be attributed to performance differences over development. Prior work has shown age-related decreases in BOLD activity during memory encoding in the right superior frontal gyrus, which mediate the relationship between age and memory performance (Tang et al., 2018). Interestingly, activity in the superior frontal gyrus during memory encoding is associated with mind-wandering and forgetting in adults (Anderson et al., 2004; Maillet & Rajah, 2016). Although detrimental to encoding, mind-wandering can be beneficial for memory consolidation (Joshi et al., 2026; Nicosia & Balota, 2024). Thus, one interpretation of this result is that memory reinstatement in the superior frontal gyrus reflects mind-wandering during rest, enhancing younger children’s memory performance. Indeed, participants who reported either spontaneously thinking about or rehearsing the memory task during the rest period exhibited better holistic episodic memory behavior, although neural memory reinstatement did not differ between those who reported consciously thinking about memories at rest or not (**Figure S11**). Alternatively, greater involvement of prefrontal regions in reinstatement in younger children could reflect higher attentional demands. Although children were not instructed to think about the memory task during the rest period, they may have recruited greater attentional resources for maintaining memory representations during the post-encoding period. In the current study, we could not tease apart conscious versus unconscious memory reactivation, which differentially affects memory performance in adults (Tal et al., 2024), nor did we have the power to investigate whether thinking about specific memories influenced behavioral performance or reinstatement for that particular memory. Future work probing children’s thoughts during awake consolidation periods will be important for better understanding mechanisms of memory reinstatement and how they may change over development.

While the superior frontal gyrus age effect was found in a whole-brain analysis, the mPFC age effect only emerged when specifically interrogated using an ROI approach. We speculated that this negative age effect could reflect increasing localization of pattern reinstatement over development (Durston et al., 2006), such that while younger children engaged a large mPFC region for pattern reinstatement, fewer voxels were recruited by older children for this same function. In a post-hoc exploratory visualization of pattern reinstatement in younger and older children, this seemed to be the case; however, caution is warranted given the exploratory nature of this analysis. In general, given the small sample size of the current study, all age analyses should be replicated in larger samples.

### Hippocampal activity during encoding relates to holistic episodic memory in development

We innovate on current models of developing memory function by focusing on holistic recall as a measure of event recollection. We predicted that hippocampal activity during encoding would relate to the successful recall of item-location-context associations given the need to pattern separate related events. Hippocampal involvement in our episodic memory task is consistent with and extends prior work in this age range (Ghetti et al., 2010; Sastre et al., 2016; Geng et al., 2019; Tang et al., 2021), including studies relating hippocampal structural volume to episodic memory behavior (DeMaster et al., 2014; Keresztes et al., 2017; Schlichting et al., 2017; Botdorf et al., 2022; Canada et al., 2019). We also found that hippocampal activity during encoding was modulated by shared scene context, such that there was less activity when youth viewed scenes that would ultimately be paired with more items. Comparable subsequent memory effects across interference trial types suggest that this effect likely reflects repetition suppression, as scenes paired with more items were repeated more often throughout the task. Repetition suppression has been documented in the adult hippocampus, particularly for scenes (Kim, 2017), and theorized to reflect the bias of memory systems to register novel information for encoding (Tulving & Kroll, 1995); here, we show such computations extend to a developmental population.

Beyond the hippocampus, we found greater activation during the encoding of remembered versus forgotten associations in regions previously associated with subsequent memory effects in adults: the lateral occipital cortex (Kim et al., 2010), supramarginal gyrus (Uncapher & Wagner, 2009), and left inferior frontal gyrus (Kirchhoff et al., 2000; Davachi et al., 2001; Staresina & Davachi, 2006; Kim, 2011). Interestingly, there were no age effects, suggesting that similar to the emergence of holistic episodic memory behavior (Ngo, Horner, et al., 2019; Ngo, Lin, et al., 2019; James et al., 2021), neural mechanisms of holistic episodic memory encoding appear rather adultlike by childhood and adolescence.

### Concluding thoughts

The current work provides novel insights into episodic memory development by investigating spontaneous reinstatement of episodic memories during an awake consolidation period, implicating the hippocampus during encoding and the angular gyrus and mPFC in the reinstatement of item-location-context associations that children and adolescents will later remember. Functional involvement of these regions is striking given their protracted structural development during childhood and adolescence. Age effects related to memory behavior were not found during encoding, but instead during post-encoding rest in the right superior frontal gyrus and mPFC, suggesting neurodevelopmental changes in prefrontal consolidation mechanisms during childhood and adolescence. These results underscore the importance of both encoding and consolidation as dynamic developmental processes that contribute to the maturation of episodic memory across childhood and adolescence.

## Resource availability

### Lead contact

Further information and requests for resources should be directed to and will be fulfilled by the lead contacts: Tristan Yates, and Nim Tottenham.

### Data and code availability

The code used to perform the analyses are available on GitHub (https://github.com/tristansyates/elfk_memory/tree/main). Due to privacy considerations, additional demographic information and raw fMRI data are not publicly available but may be shared by the lead contacts upon request.

## Acknowledgements

We are thankful to the families who participated in this research. This work was supported by New York State Psychiatric Institute seed funding to B.C. and N.T. T.S.Y was supported by an NIH F32 fellowship (1F32HD114417-01A1). B.C. was supported by a NARSAD Young Investigator Grant (24739). We are grateful to the members of the DAN and DCN Labs at Columbia for feedback on earlier versions of this project. We are grateful to Naeun Oh for code review.

## Author contributions

**T.S.Y.:** Conceptualization, Software, Formal analysis, Writing – Original Draft, Writing – Reviewing & Editing; **B.C.:** Conceptualization, Methodology, Investigation, Funding Acquisition, Writing – Reviewing & Editing; **L.A.S.:** Conceptualization, Methodology, Investigation, Writing – Reviewing & Editing; **M.V.:** Investigation, Writing – Reviewing & Editing; **T.C.:** Investigation, Writing – Reviewing & Editing; **K.O.S.:** Investigation, Writing – Reviewing & Editing; **L.D.:** Conceptualization, Methodology, Writing – Reviewing & Editing, Supervision; **N.T.:** Conceptualization, Methodology, Funding Acquisition, Writing – Reviewing & Editing, Supervision.

## Methods

### Experimental model and study participant details

#### Participants

Children and adolescents were invited to participate in an fMRI study between November 2015 and April 2018 (Callaghan et al., 2021; Silvers et al., 2021; Abramson et al., 2024). Participants were recruited through local birth records, local classifieds, posted flyers, and referrals. Of the 95 participants who participated in the episodic memory encoding task during fMRI, behavioral data could be analyzed from 84 participants (8 excluded for not completing the memory test, 1 excluded for encoding task timing error, 1 excluded for falling asleep during encoding, and 1 excluded for computer error during the memory test). For the analyses of the current paper, we focused on children and adolescents with typical developmental histories and therefore did not consider participants who had a history of early adversity in the form of previous institutionalization (*N* = 32) or who experienced domestic foster care followed by adoption in the United States (*N* = 3). Behavioral analyses were run on a final sample of 49 participants (age: *M* = 11.68 years, range = 6.65 – 17.61 years, 26 female). Memory encoding analyses were run on 46 participants (2 excluded for missing anatomical image, 1 excluded for excess motion during the encoding run; see *fMRI Preprocessing* section below). Of these participants, reinstatement analyses were run on 42 participants (1 excluded for missing post-encoding rest run, 3 excluded for excess motion during pre-and/or post-encoding rest).

Caregivers provided informed written consent and children/adolescents provided assent (verbal assent for children under age 7 and written assent for children over age 7). Behavioral data and analyses on hippocampal granularity were previously reported in (Callaghan et al., 2021). All procedures were approved by the Institutional Review Board at Columbia University.

## Method details

### Data acquisition

MRI data were acquired at the New York State Psychiatric Institute on a 3T General Electric SIGNA MRI scanner with a NOVA 32-channel coil. Functional data were collected using a T2*-weighted gradient-EPI sequence collected at an oblique angle (TR = 2000 s, TE = 30 ms, flip angle = 75°, FOV = 200 mm, voxel size = 3.125 × 3.125 × 4 mm, interleaved slice acquisition, 20° oblique angle). Structural images (T1-weighted; SPGR) were collected for anatomical alignment.

### Procedure

Prior to the MRI scan session, participants came in for a behavioral visit in which they completed several questionnaires and cognitive tasks that will not be described here. On the day of the MRI scan, participants were first given task instructions and an opportunity to practice responding to the tasks they would perform in the scanner. For the memory task, participants were explicitly told that they would be playing a ‘hide and seek game’ in which they would need to remember where toys and ‘friends’ (faces of unknown children) were hiding in different scenes. Then, participants entered the scanner and performed an initial resting state scan in which they stared at a fixation cross for 5 minutes (pre-encoding rest), followed by the memory encoding task (described below), and finally a second resting state scan (post-encoding rest; see **Figure 1A**). During the resting state scans, participants were not given any instructions about what to think about. A high-resolution anatomical scan was then collected while participants viewed movies. Finally, additional tasks were run in the scanner that are beyond the scope of the current manuscript. Outside of the scanner, participants performed an immediate memory test and a debrief questionnaire. Approximately two weeks later, some participants completed an identical, offline delayed memory test in their homes, which is not discussed here.

### Memory encoding fMRI task

Participants performed an associative memory encoding task during a single fMRI run (described in detail in Callaghan et al., 2021). The task was presented using E-Prime version 2.0 software (Psychology Software Systems) and was approximately 7 minutes in length. On each trial of the task, a cartoonized scene appeared for 500 ms before an item (either a cartoon toy or a child’s face) appeared in a location on the scene for an additional 2500 ms (**Figure 1A**). Each encoding trial was followed by a jittered intertrial interval in which left and right arrows appeared on the screen (average jitter = 8.5 s; range: 4.2 to 16.8 s), before a one-second fixation cross appeared prior to the next trial. Participants were asked to press a button on the button box when they saw an arrow during the baseline to maintain engagement during an otherwise passive viewing task. Participants saw a total of 20 unique item-location-context pairs, which consisted of 7 scenes that were each paired with either 2 or 4 items. These item-location-context pairs were presented either once or twice during the memory encoding task, resulting in 33 total encoding trials. For the current analyses, we do not examine the effects of repetition in order to increase power. Participants saw the memory encoding trials in one of two fixed orders.

### Immediate memory test

After exiting the scanner, participants performed a self-paced memory test without feedback. The memory test was presented in E-Prime version 2.0 in a randomized order. Children responded to questions verbally or via pointing while a researcher recorded their answer, and adolescents made their own responses while a researcher sat beside them. The test phase consisted of two parts: an item recognition memory test and an associative memory test. During the item recognition memory test, participants were asked to indicate whether an item was ‘old,’ ‘new,’ or if they didn’t know. Participants completed 40 recognition memory test trials, consisting of 20 old items and 20 new foils. Foils were chosen to closely match one of the items seen at encoding (e.g., if a toy truck was an old item, a toy motorcycle was the foil). Results from the recognition memory task are beyond the scope of the current paper. Then, participants completed an associative memory test on each of the 20 item-location-context association pairs (**Figure 1B**). First, participants saw a scene with three objects or faces (all of which had been previously seen during encoding) and were asked to pick which of the toys/faces had been paired with the scene (‘which was hiding here?’). Then (without feedback), participants saw circles with numbers inside (in yellow) overlaid on the scene and were asked to choose the number which corresponded to the location in which the toy/face had appeared on the scene (‘where was it hiding?’). All test locations were potential locations for an item seen at encoding — in other words, for scenes that had been paired with two different items, participants had the option of two different locations, and for scenes that that had been paired with four items, participants had the option of four different locations. We defined ‘holistic episodic memory’ as participants’ ability to correctly retrieve both the item that was paired with the scene and its location on the scene.

On average, the memory test occurred 56.79 minutes after encoding (range = 19.43 to 83.45 minutes; SD = 12.70 minutes; timing delay information missing from *N* = 5 participants). We did not find a significant main effect of delay length on memory performance (*β* = −0.0038, *p* = .660), nor an interaction between delay length and age (*β* = −0.006, *p* = .898). Thus, we do not consider the delay between encoding and test further, although note the difference in delays across participants as a potential limitation.

## Quantification and statistical analysis

### fMRIPrep preprocessing

Preprocessing of the MRI data was performed using fMRIPrep v. 23.1.3 (Esteban et al., 2019). The full boilerplate text generated by fMRIPrep, which is covered by a CC0 license and intended to be copied verbatim (see https://fmriprep.org/en/20.2.0/citing.html), is located in the **Supplementary Text**.

In brief, the anatomical T1-weighted image was corrected for intensity non-uniformity and skull-stripped (ANTs; Avants et al., 2008) before brain tissue segmentation of cerebrospinal fluid (CSF), white-matter (WM) and gray-matter (GM) using fast with FSL (Zhang et al., 2001). Volume-based spatial normalization to standard space (MNI152NLin6Asym) was performed using nonlinear registration (ANTs; Avants et al., 2008). For each functional run that was available per participant (up to three runs: pre-encoding rest, memory encoding, and post-encoding rest), a reference volume and its skull-stripped version were created. Head-motion parameters with respect to the reference (transformation matrices, and six corresponding rotation and translation parameters) were estimated using mcflirt from FSL. Functional runs were slice-time corrected using 3dTshift from AFNI (Cox & Hyde, 1997) and then resampled to native space after applying the transforms correcting for head motion. Functional runs were registered to the participant’s T1 anatomical image using mri_coreg (FreeSurfer; Fischl, 2012) and flirt from FSL with 6 degrees of freedom. Finally, functional runs were registered to standard space (MNI152NLin6Asym) using ANTs (Avants et al., 2008). For subsequent analyses, we used functional runs registered to standard space in native voxel resolution (i.e., 3.125 x 3.125 x 4mm voxels). Functional runs were also spatially smoothed (5 mm) and high-pass filtered (100 s) before being fit to general linear models (GLMs). For the pattern reinstatement analyses, we did not use spatial smoothing in order to maintain fine-grained response patterns (Weaverdyck et al., 2020). In all subsequent analyses, timepoints in which the framewise displacement (FD) exceeded 0.9 mm were treated as motion outliers and runs with more than 30% of motion outliers were excluded from further analysis (1 memory encoding run and 3 rest runs). For the included runs, the proportion of motion outliers was low (pre-encoding rest: *M* = 1.2%, *SD* = 3.0%; memory encoding: *M* = 4.5%, *SD* = 7.3%; post-encoding rest: *M* = 2.3%, *SD* = 3.6%), as was the average framewise displacement (pre-encoding rest: *M* = 0.15, *SD* = 0.10, memory encoding: *M* = 0.22, *SD* = 0.21; post-encoding rest: *M* = 0.18, *SD* = 0.12). The relationship between age and the average FD across participants for each of the three runs is visualized in **Figure S12**.

### Holistic episodic memory behavior

Our main analyses focused on holistic episodic memory — that is, whether participants remembered both the association between an item and its scene context and the location at which the item appeared on the scene. As in prior work (Callaghan et al., 2021), we did not require that children correctly identify an item as ‘old’ in the recognition memory task for them to receive credit on the associative memory test for that item. For our main behavioral analyses, holistic episodic memory performance was calculated as the proportion of correct trials out of 20 total trials. Chance performance varied depending on whether a scene was paired with two items (16.67%) or with four items (8.33%), and so we assessed whether holistic episodic memory was significantly above the average chance level (12.5%) using two-sided, one-sample *t*-tests. We additionally tested whether behavior was above chance for trials in which the scene was paired with two items (N = 8 trials; ‘low interference’) or four items (N = 12 trials; ‘high interference’) with the appropriate chance levels. Age effects were tested using Pearson correlation.

### Anatomically defined regions of interest

For our neural analyses, we primarily took a whole-brain approach, but also examined regions of interest (ROIs) used in prior research on reinstatement in adults (Yu et al., 2024). We combined parcels from the Schaefer atlas (Schaefer et al., 2018) to create ROIs for ventral temporal cortex, retrosplenial cortex, and medial prefrontal cortex (see **Table S1**). We additionally created an ROI for the angular gyrus given recent work showing its involvement in item-level reinstatement and memory precision (Hou et al., 2025). The bilateral hippocampus was defined using the Harvard-Oxford subcortical atlas (Jenkinson et al., 2012) (25% max probability). For follow-up exploratory analyses of age and trial type on encoding and reinstatement, we created empirically derived ROIs from our main whole-brain analyses.

### Univariate encoding analysis

To examine how neural activity during encoding relates to holistic episodic memory performance during development, we conducted a subsequent memory analysis (Brewer et al., 1998; Wagner et al., 1998). This within-participant analysis identifies brain regions that are more active during the encoding of trials that would later be remembered or forgotten at the individual participant level. Each encoding trial was modeled using a boxcar for its duration (3 s) and convolved with a double-gamma hemodynamic response function. Trials were then back-coded as either subsequently remembered or subsequently forgotten based on that participant’s holistic episodic memory at test. As in our behavioral memory analyses, participants needed to remember both the pairing of an item with its scene context and the location that it appeared on the scene for a trial to be considered remembered (otherwise it was considered ‘forgotten’). For item-location-context pairings that were seen twice during encoding, we treated each viewing as an independent trial because we did not have power to examine the effects of repetition. We fit GLMs to preprocessed BOLD activity with one regressor for all subsequently remembered trials and one regressor for all subsequently forgotten trials using FEAT with FSL. The temporal derivatives of these regressors were included in the design matrix to achieve a better fit to the data, and pre-whitening was employed to account for temporal autocorrelation and improve statistical efficiency. Additional regressors of no interest included 12 motion parameters (3 translation, 3 rotation, and their temporal derivatives) and the mean signal in CSF and WM tissues. Time points excluded for high motion (> 0.9 mm framewise displacement) were scrubbed using additional regressors of no interest. For each participant, we extracted the whole-brain *z*-statistic volume for the contrast of subsequently remembered encoding trials greater than subsequently forgotten encoding trials. Analyses were run at native voxel resolution within participant but resampled to standard space voxel resolution (2mm isometric) for whole-brain group statistics.

As a follow-up exploratory analysis, we used a similar approach to model the influence of shared scene context on encoding activity. This analysis was conducted in two different ways. First, we followed the same general procedure as above, except that the two main regressors were ‘low interference trials’ (i.e., trials comprising scenes that would be paired with two items) and ‘high interference trials’ (i.e., trials comprising scenes that would be paired with four items). The contrast of interest was high versus low interference trials. Second, we ran a subsequent memory analysis contrasting remembered and forgotten trials separately for low and high interference trials. This analysis separates a relatively small number of trials into 4 regressors (remembered/forgotten x low/high interference), with 6 participants dropped from the analysis for not having at least one trial in each of the 4 regressors.

### Multivariate pattern reinstatement analysis

We next examined how reinstatement of previously encoded episodes during post-encoding rest relates to holistic episodic memory performance. We conducted subsequent memory reinstatement analyses inspired by previous work investigating the reinstatement of encoded content during rest runs (Staresina et al., 2013; Schapiro et al., 2018; Jimenez & Meyer, 2024). Specifically, after preprocessing of the rest runs, we regressed out 12 motion parameters (3 translation, 3 rotation, and their temporal derivatives), the mean signal in CSF and WM tissues, and time points excluded for high motion (> 0.9 mm framewise displacement) for each rest run of each participant. These preprocessed volumes were then *z*-scored over time. For encoding runs, we ran a GLM that modeled each individual encoding trial as a single regressor (33 total regressors), along with the same regressors of no interest used for the rest runs. We extracted the *z*-statistic images from these GLMs as our measure of encoding activity. Analyses were run at native voxel resolution within participant but resampled to standard space voxel resolution (2mm isometric) for whole-brain group statistics.

We conducted a cubical searchlight analysis (7 x 7 x 7 voxels) across the whole brain using BrainIAK (Kumar et al., 2021). In each searchlight, we first correlated the pattern of activity for each encoding trial with the pattern of activity at each TR in the pre-encoding rest run and the post-encoding rest run (150 TRs each; **Figure 3A**). These encoding trial by rest TR correlation matrices were then split into trials that would be subsequently remembered versus subsequently forgotten. Correlation values were summed within memory type (remembered versus forgotten) and rest run (pre-encoding and post-encoding). Our reinstatement metric was calculated as the difference between these summed correlation values for remembered minus forgotten trials in the post-encoding rest run after accounting for the pre-encoding rest run (i.e., [RememberedPost – Remembered_Pre_] – [Forgotten_Post_ – Forgotten_Pre_]), with the logic that there should not be reinstatement prior to memory encoding. Reinstatement values were compared to a chance level of zero.

For this within-participant analysis, we opted not to threshold correlation values in the encoding-rest correlation matrices prior to summing the correlation values. Prior work examining individual differences in memory reinstatement defined potential reinstatement events as those timepoints at which the correlations between encoding and rest were greater than 1.5 standard deviations from the mean within a given rest run (Schapiro et al., 2018; Jimenez & Meyer, 2024); here, we found that such an approach diminished power for looking at the contrast of subsequently remembered vs forgotten trials within participant and that changes in the threshold value gave different results. To validate our threshold-free approach, we re-ran the reinstatement analyses after randomly shuffling remembered and forgotten trails labels 1000 times, recalculating the reinstatement metric on each iteration. Across regions, these permuted values were not different from chance (**Figure S6**).

Finally, we ran two exploratory follow up analyses to examine reinstatement of low and high interference trials. The first analysis examined whether high interference trials were reinstated more than low interference trials during the post-encoding rest run after correcting for the pre-encoding rest run (i.e., [High_Post_ – High_Pre_] – [Low_Post_ – Low_Pre_]). The second analysis examined whether there was greater reinstatement of remembered versus forgotten trials in the post-encoding rest run separately for low and high interference trials. For this latter analysis, participants who did not have at least one trial that was remembered and one trial that was forgotten were dropped from the analysis (low interference: *N* = 2 dropped, high interference: *N* = 6 dropped).

### Statistical analysis

For testing significance across the whole-brain, we used FSL’s randomise tool to conduct non-parametric statistics (Winkler et al., 2014). This consisted of a sign test, in which the signs of the data were randomly flipped across 1000 iterations and *p*-values were calculated based on this permuted distribution. We performed multiple-comparisons correction on the whole-brain *t*-statistic maps from randomise using FSL’s cluster tool (cluster-forming threshold: *p* < .05, *z*-statistic threshold = 2.41, smoothness estimate from the *z*-statistic map).

For testing significance within ROIs for both encoding and reinstatement analyses, we examined whether a given contrast of interest (e.g., remembered versus forgotten encoding trials) differed from zero using two-sided, one-sample *t*-tests. Age effects were tested using Pearson correlation.

## Supplementary Materials

### Supplementary Text

#### Anatomical data preprocessing

A total of 1 T1-weighted (T1w) images were found within the input BIDS dataset. The T1-weighted (T1w) image was corrected for intensity non-uniformity (INU) with N4BiasFieldCorrection (Tustison et al., 2010), distributed with ANTs (2.4.4; Avants et al., 2008), RRID:SCR_004757), and used as T1w-reference throughout the workflow. The T1w-reference was then skull-stripped with a Nipype implementation of the antsBrainExtraction.sh workflow (from ANTs), using OASIS30ANTs as target template. Brain tissue segmentation of cerebrospinal fluid (CSF), white-matter (WM) and gray-matter (GM) was performed on the brain-extracted T1w using fast (FSL 6.0.6.2, RRID:SCR_002823, Zhang et al., 2001). Volume-based spatial normalization to two standard spaces (MNI152NLin6Asym, MNI152NLin2009cAsym) was performed through nonlinear registration with antsRegistration (ANTs 2.4.4), using brain-extracted versions of both T1w reference and the T1w template. The following templates were selected for spatial normalization and accessed with TemplateFlow (23.0.0, Ciric et al., 2022): FSL’s MNI ICBM 152 non-linear 6th Generation Asymmetric Average Brain Stereotaxic Registration Model [Evans et al., (2012), RRID:SCR_002823; TemplateFlow ID: MNI152NLin6Asym], ICBM 152 Nonlinear Asymmetrical template version 2009c [Fonov et al., (2009), RRID:SCR_008796; TemplateFlow ID: MNI152NLin2009cAsym].

#### Functional data preprocessing

For each of the 3 BOLD runs found per subject (across all tasks and sessions), the following preprocessing was performed. First, a reference volume and its skull-stripped version were generated using a custom methodology of fMRIPrep. Head-motion parameters with respect to the BOLD reference (transformation matrices, and six corresponding rotation and translation parameters) are estimated before any spatiotemporal filtering using mcflirt (FSL 6.0.6.2, Jenkinson et al., 2012). BOLD runs were slice-time corrected to 1s (0.5 of slice acquisition range 0s-2s) using 3dTshift from AFNI (Cox & Hyde, 1997, RRID:SCR_005927). The BOLD time-series (including slice-timing correction when applied) were resampled onto their original, native space by applying the transforms to correct for head-motion. These resampled BOLD time-series will be referred to as preprocessed BOLD in original space, or just preprocessed BOLD. The BOLD reference was then co-registered to the T1w reference using mri_coreg (FreeSurfer) followed by flirt (FSL, Jenkinson & Smith, 2001) with the boundary-based registration (Greve & Fischl, 2009) cost-function. Co-registration was configured with six degrees of freedom.

Several confounding time-series were calculated based on the preprocessed BOLD: framewise displacement (FD), DVARS and three region-wise global signals. FD was computed using two formulations following Power (absolute sum of relative motions, Power et al., 2014) and Jenkinson (relative root mean square displacement between affines, Jenkinson et al., 2002). FD and DVARS are calculated for each functional run, both using their implementations in Nipype (following the definitions by Power et al., 2014). The three global signals are extracted within the CSF, the WM, and the whole-brain masks. Additionally, a set of physiological regressors were extracted to allow for component-based noise correction (CompCor, Behzadi et al., 2007). Principal components are estimated after high-pass filtering the preprocessed BOLD time-series (using a discrete cosine filter with 128s cut-off) for the two CompCor variants: temporal (tCompCor) and anatomical (aCompCor). tCompCor components are then calculated from the top 2% variable voxels within the brain mask. For aCompCor, three probabilistic masks (CSF, WM and combined CSF+WM) are generated in anatomical space. The implementation differs from that of Behzadi et al. in that instead of eroding the masks by 2 pixels on BOLD space, a mask of pixels that likely contain a volume fraction of GM is subtracted from the aCompCor masks. This mask is obtained by thresholding the corresponding partial volume map at 0.05, and it ensures components are not extracted from voxels containing a minimal fraction of GM. Finally, these masks are resampled into BOLD space and binarized by thresholding at 0.99 (as in the original implementation). Components are also calculated separately within the WM and CSF masks. For each CompCor decomposition, the k components with the largest singular values are retained, such that the retained components’ time series are sufficient to explain 50 percent of variance across the nuisance mask (CSF, WM, combined, or temporal). The remaining components are dropped from consideration. The head-motion estimates calculated in the correction step were also placed within the corresponding confounds file. The confound time series derived from head motion estimates and global signals were expanded with the inclusion of temporal derivatives and quadratic terms for each (Satterthwaite et al., 2013). Frames that exceeded a threshold of 0.9 mm FD or 1.5 standardized DVARS were annotated as motion outliers. Additional nuisance timeseries are calculated by means of principal components analysis of the signal found within a thin band (crown) of voxels around the edge of the brain, as proposed by (Patriat et al., 2017). The BOLD time-series were resampled into standard space, generating a preprocessed BOLD run in MNI152NLin6Asym space. First, a reference volume and its skull-stripped version were generated using a custom methodology of fMRIPrep. All resamplings can be performed with a single interpolation step by composing all the pertinent transformations (i.e. head-motion transform matrices, susceptibility distortion correction when available, and co-registrations to anatomical and output spaces). Gridded (volumetric) resamplings were performed using antsApplyTransforms (ANTs), configured with Lanczos interpolation to minimize the smoothing effects of other kernels (Lanczos 1964). Non-gridded (surface) resamplings were performed using mri_vol2surf (FreeSurfer).

Many internal operations of fMRIPrep use Nilearn 0.10.1 (Abraham et al., 2014, RRID:SCR_001362), mostly within the functional processing workflow. For more details of the pipeline, see the section corresponding to workflows in fMRIPrep’s documentation.

## Supplementary Tables

**Table S1.**
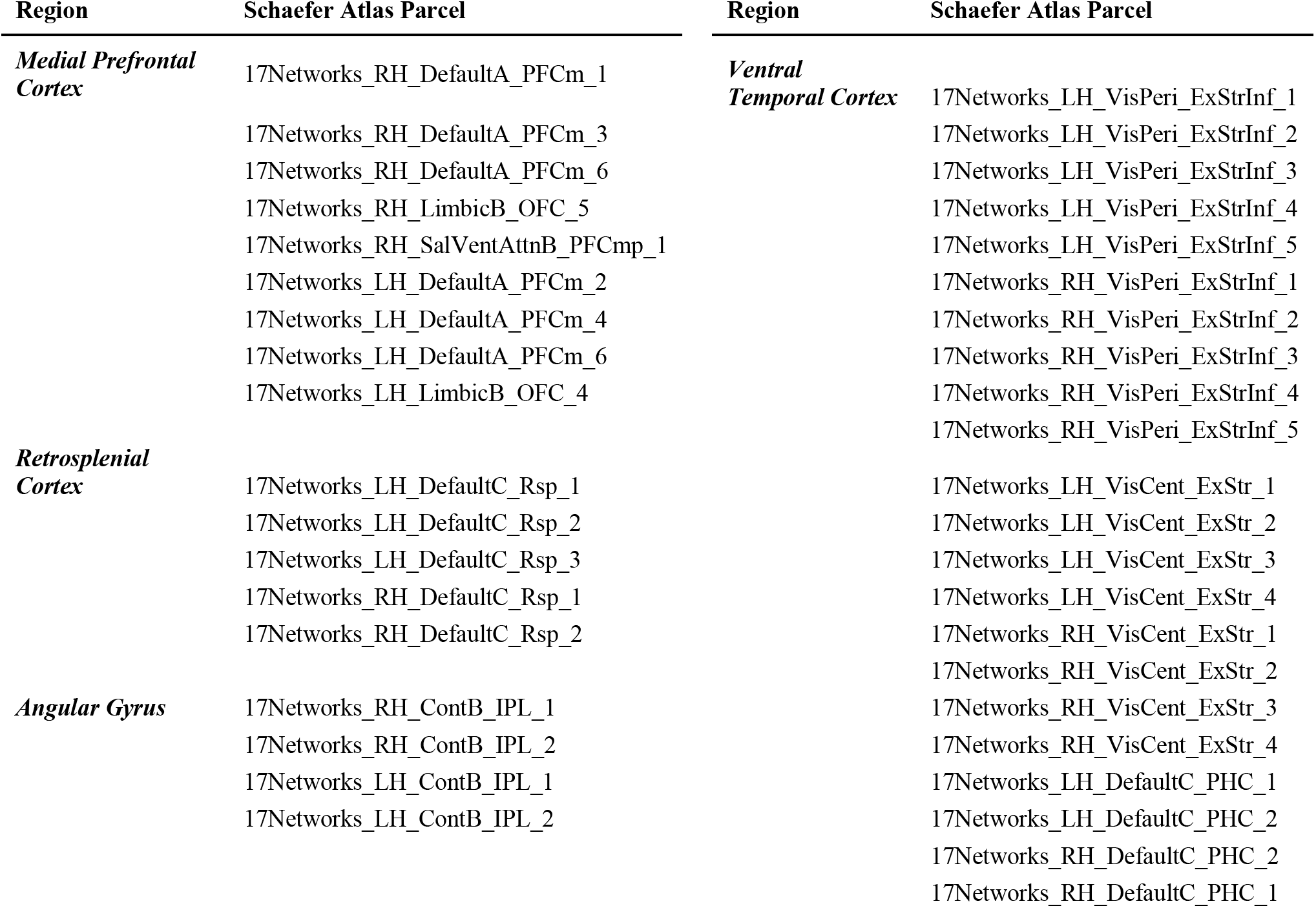
Parcels comprising ROIs used in prior adult work. Labels for the parcels from the Schaefer atlas (Number of parcels = 400, Number of Yeo Networks =17, Resolution = 2mm) (Schaefer et al., 2018) that were used to create the cortical ROIs. Note that the hippocampus is not listed as it was created using the Harvard-Oxford atlas.

**Table S2.** Memory encoding in the whole brain. Regions showing a significant effect during encoding in a whole brain voxel-wise analysis following multiple comparisons correction (cluster-defining threshold: *z* > 2.41, *p* < .05). Coord = coordinate, COG = center of gravity, Rem = remembered, Forg = forgotten.

| Univariate Activity during Encoding |  |  |  |  |  |  |  |  |  |
| --- | --- | --- | --- | --- | --- | --- | --- | --- | --- |
| Contrast | Region | Cluster Size (mm<sup>3</sup>) | Peak Coord X | Peak Coord Y | Peak Coord Z | Peak Z-value | COG Coord X | COG Coord Y | COG Coord Z |
| ***Rem > Forg*** | Right Anterior Hippocampus | 2760 | 26 | -14 | -20 | 4.93 | 38.9 | -33.3 | -12.6 |
|  | Left Lateral Occipital Cortex | 2530 | -50 | -70 | -8 | 4.93 | -51.2 | -56.7 | -9.1 |
|  | Left Inferior Frontal Gyrus | 1428 | -40 | 6 | 24 | 4.57 | -44.7 | 5.8 | 32.7 |
|  | Left Supramarginal Gyrus | 1124 | -64 | -34 | 36 | 4.38 | -55.5 | -32.7 | 41.1 |
|  | Right Supramarginal Gyrus | 1032 | 54 | -28 | 50 | 4.76 | 48.7 | -35.2 | 51.0 |

**Table S3.** Pattern reinstatement at rest in the whole brain. Regions showing a significant effect during post-encoding rest in a whole brain voxel-wise analysis following multiple comparisons correction (cluster-defining threshold: *z* > 2.41, *p* < .05). Coord = coordinate, COG = center of gravity, Rem = remembered, Forg = forgotten.

| Pattern Reinstatement at Rest |  |  |  |  |  |  |  |  |  |
| --- | --- | --- | --- | --- | --- | --- | --- | --- | --- |
| Contrast | Region | Cluster Size (mm3) | Peak Coord X | Peak Coord Y | Peak Coord Z | Peak Z-value | COG Coord X | COG Coord Y | COG Coord Z |
| *Rem > Forg* | Medial Prefrontal Cortex | 4570 | -28 | 36 | 34 | 3.76 | -17.4 | 35.2 | 29.4 |
|  | Left Angular Gyrus | 4438 | -50 | -62 | 6 | 4.93 | -50.5 | -60.4 | 28.9 |
|  | Right Angular Gyrus | 2380 | 48 | -74 | 36 | 3.80 | 44.3 | -55.8 | 44.2 |
|  | Left Precentral Gyrus | 1712 | -50 | 4 | 40 | 3.97 | -44.3 | -0.2 | 45.5 |

## Supplementary Figures

**Figure S1.**
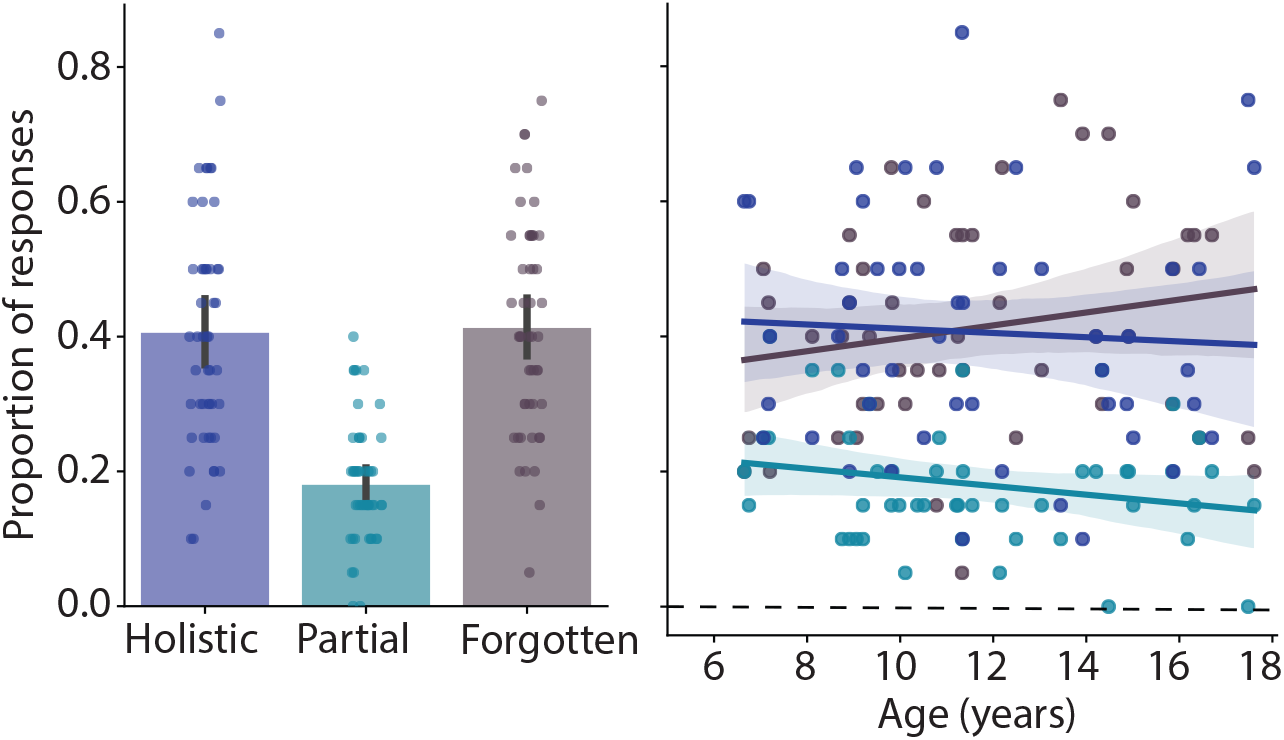
Different behavioral responses during memory test. Behavior during the associative memory test could be broken down into three categories: holistic memory (both item-context and item-location correct), partial memory (item-context correct and item-location incorrect), and no memory (both item-context and item-location questions incorrect, *or* only item-location question correct). There were few trials (average < 20%) in which children had only partial memory, with no relationship to age. Dots represent individual participant data and shaded error bars and regions reflect the 95% confidence interval from bootstrap resampling.

**Figure S2.**
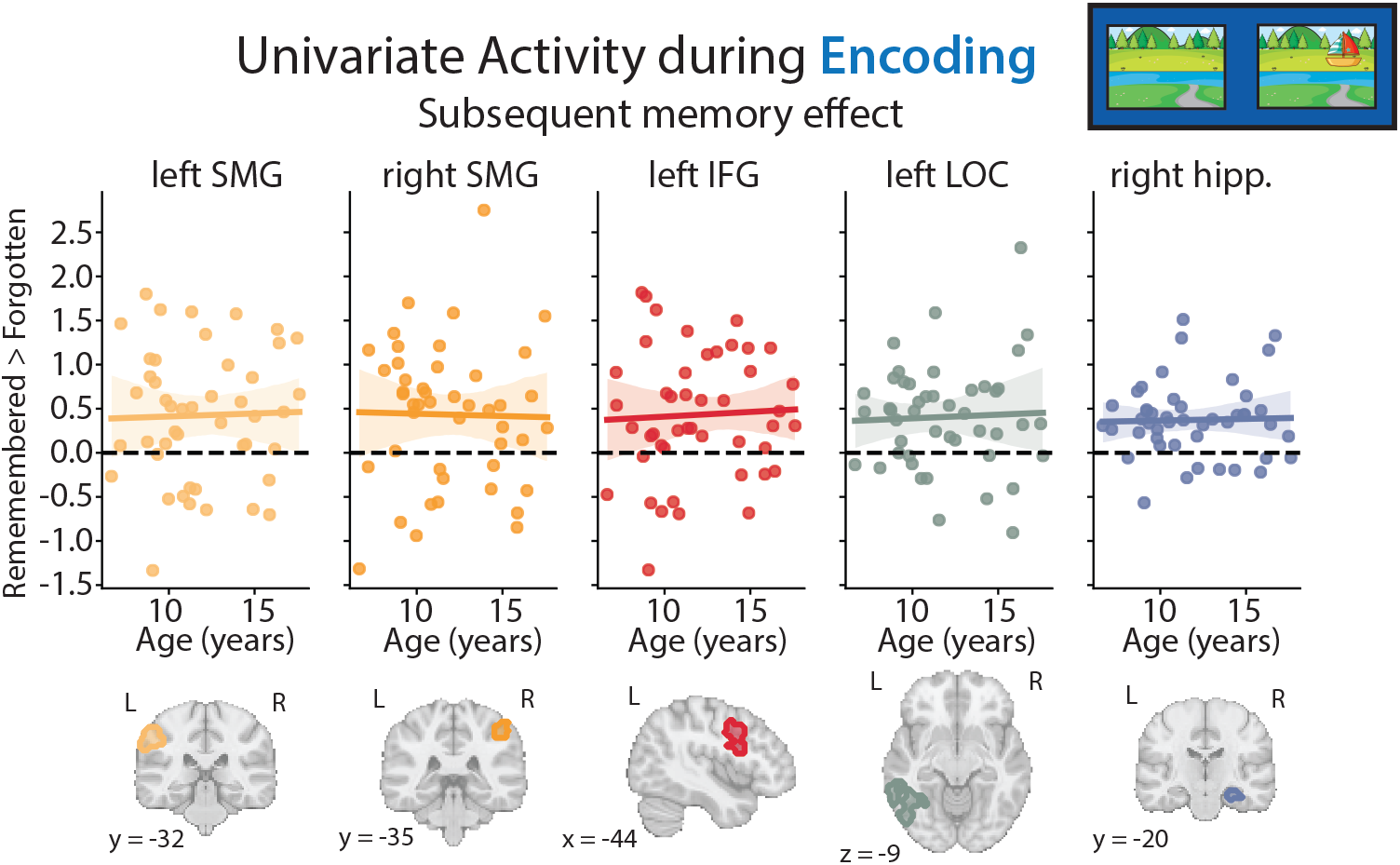
Effects of age on neural mechanisms of episodic memory encoding. Relationship between age and the contrast of subsequently remembered versus subsequently forgotten trials during memory encoding in regions significant in the whole brain (Figure 2**; Table S2)**. There was no effect of age in any of the regions (all *p*s >.768). Dots represent individual participant data and shaded error regions reflect the 95% confidence interval from bootstrap resampling. SMG = supramarginal gyrus, IFG = inferior frontal gyrus, LOC = lateral occipital cortex, hipp. = hippocampus.

**Figure S3.**
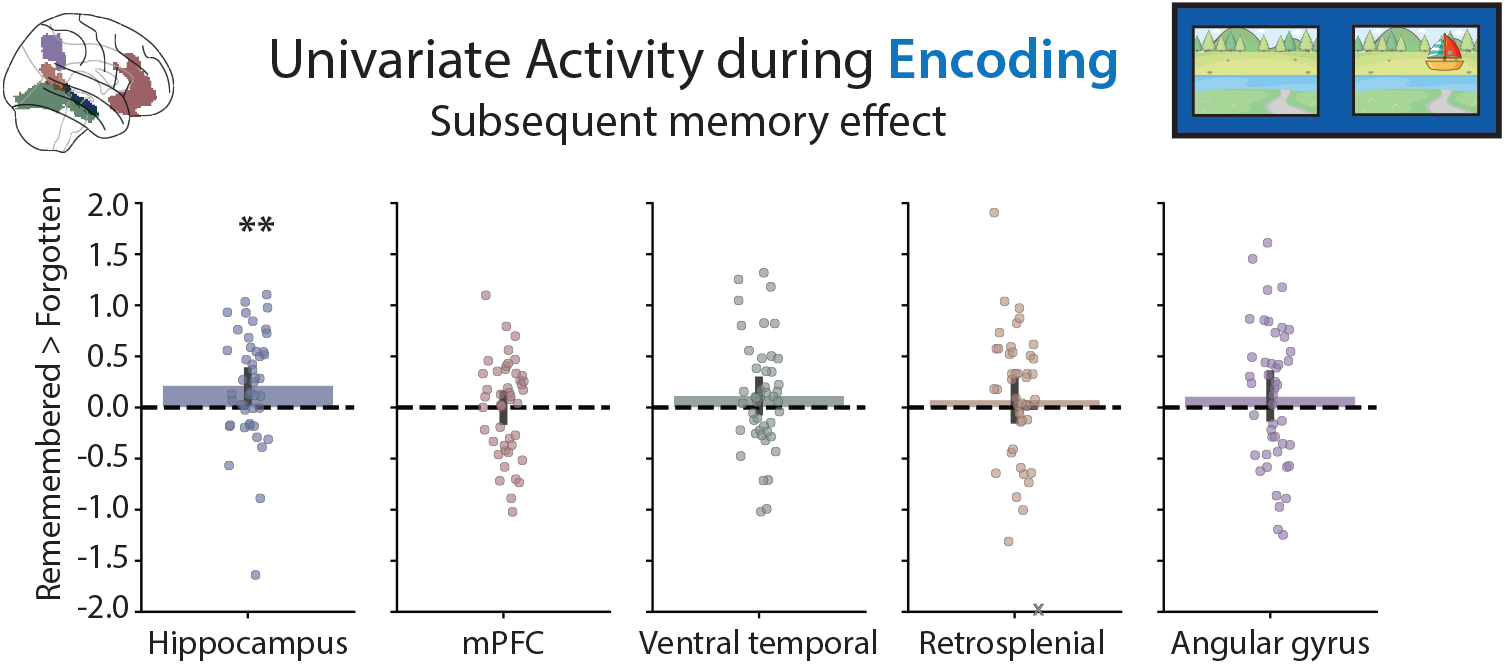
Neural mechanisms of memory encoding in anatomically defined ROIs. Contrast of subsequently remembered versus subsequently forgotten trials during memory encoding in anatomically defined ROIs from the adult literature. ROIs are visualized on a glass brain in the top left corner. Youth showed significantly greater activation in the bilateral hippocampus during the encoding of trials that would later be remembered versus forgotten (*M* = 0.22, *t*(45) = 2.75, *p* = .009, *d* = 0.41); no other ROI showed a significant subsequent memory effect (mPFC: *M* = 0.00, *t*(45) = 0.01, *p* = .995, *d* = 0.00; ventral temporal: *M* = 0.11, *t*(45) = 1.42, *p* = .164, *d* = 0.21; retrosplenial: *M* = 0.07, *t*(45) = 0.69, *p* = .496, *d* = 0.10; angular gyrus: *M* = 0.11, *t*(45) = 1.07, *p* = .289, *d* = 0.16). There was no effect of age on the subsequent memory contrast in any of these anatomically defined ROIs (all *p*s > .179). Dots represent individual participant data and error bars reflect the 95% confidence interval from bootstrap resampling. One participant with a value outside of the plotting range for the retrosplenial cortex is marked with an X on the negative edge of the y-axis. ** *p* < .01.

**Figure S4.**
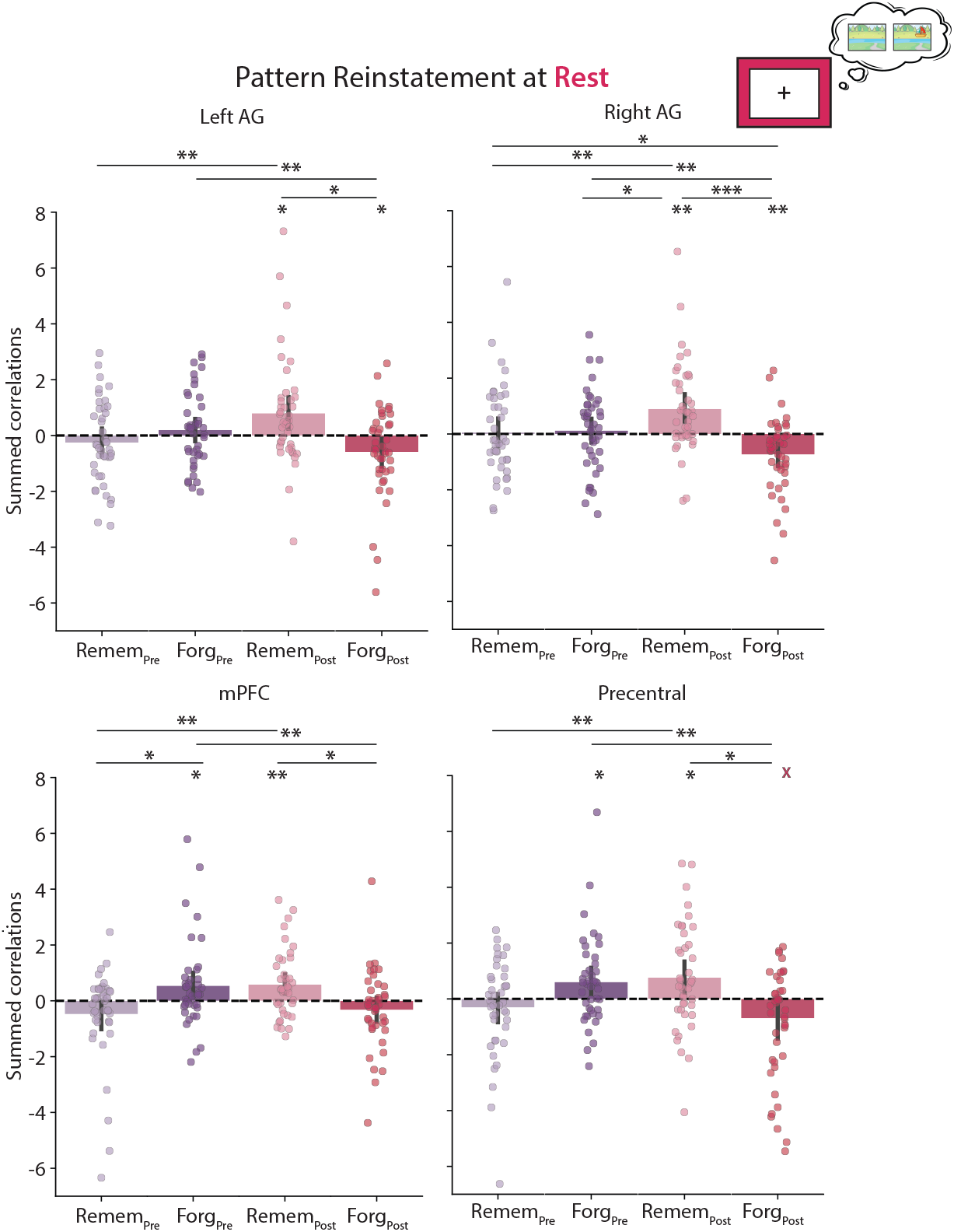
Breakdown of spontaneous pattern reinstatement during rest in empirical ROIs. The sums of the thresholded correlations for each trial type (remembered or forgotten) and rest run (pre-encoding or post-encoding) are visualized separately for the four regions that were significant in the whole-brain searchlight analysis (Figure 3**; Table S3**). To create the reinstatement *z*-statistics used in the main analysis, we subtracted the summed correlation for remembered minus forgotten trials during post-encoding rest after correcting for pre-encoding rest. Dots represent individual subject data and error bars reflect the 95% confidence interval from bootstrap resampling. *** *p* < .001, ** *p* < .01, * *p* < .05. mPFC = medial prefrontal gyrus, AG = angular gyrus, Remem = remembered, Forg = forgotten.

**Figure S5.**
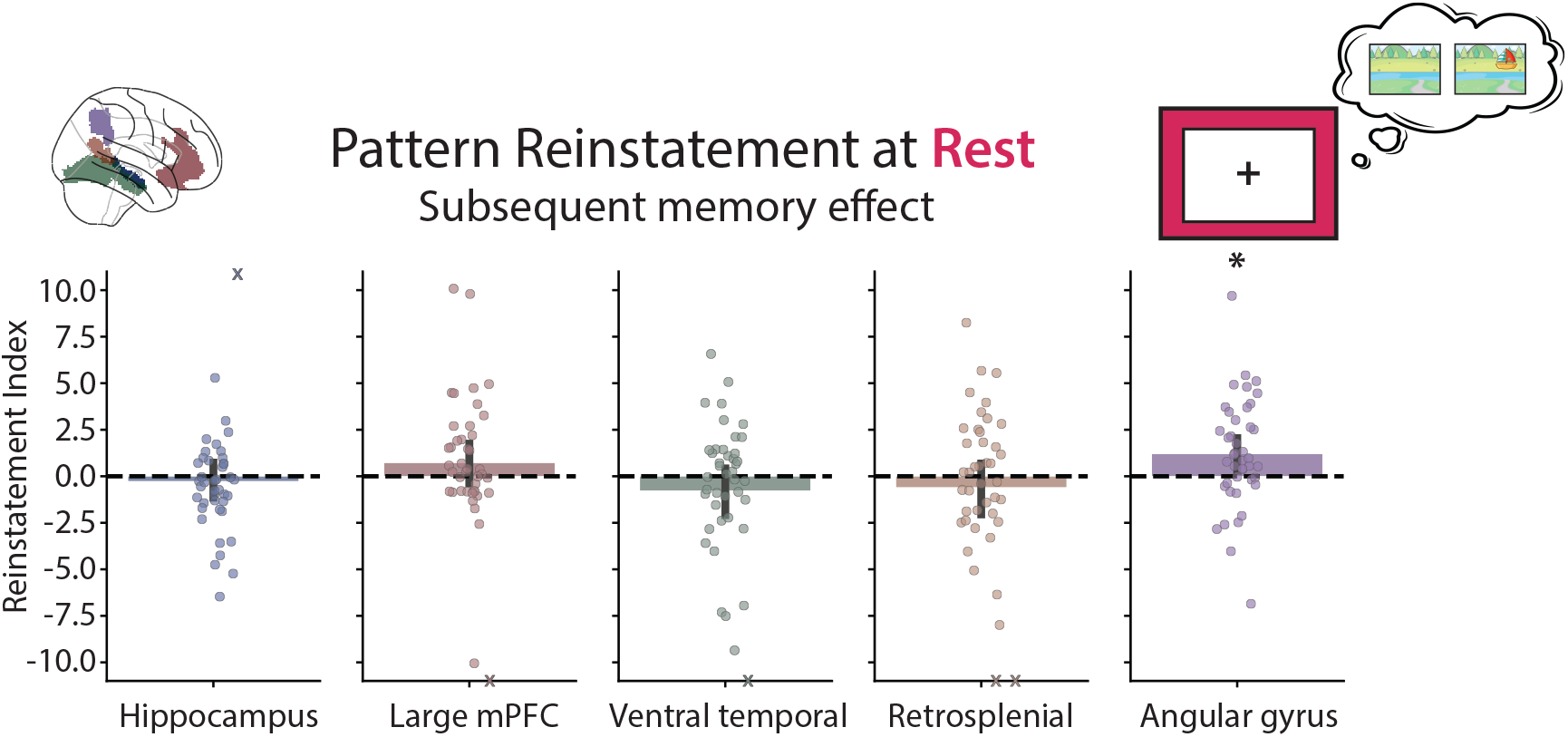
Spontaneous pattern reinstatement in anatomically defined ROIs. Spontaneous memory reinstatement results showing the contrast of subsequently remembered versus subsequently forgotten trials during post-encoding rest in anatomically defined ROIs from the adult literature. ROIs are visualized on a glass brain in the top left corner. During the post-encoding rest scan, youth showed significantly more pattern reinstatement of trials that would later be remembered versus forgotten, controlling for the pre-encoding rest scan, in the angular gyrus (*M* = 1.20, *t*(41) = 2.65, *p* = .011, *d* = 0.41) but not in the other ROIs (hippocampus: *M* = −0.26, *t*(41) = −0.55, *p* = .587, *d* = −0.09; mPFC: *M* = 0.72, *t*(41) = 1.26, *p* = .216, *d* = 0.20; ventral temporal: *M* = −0.77, *t*(41) = −1.20, *p* = .238, *d* = −0.19; retrosplenial: *M* = −0.60, *t*(41) = −0.84, *p* = .408, *d* = −0.13). There was no effect of age on subsequent memory reinstatement in any of these anatomically defined ROIs (all *p*s > .400). Dots represent individual participant data and error bars reflect the 95% confidence interval from bootstrap resampling. Participants with values outside of the plotting range are marked with Xs on the positive or negative edge of the y-axis. * *p* < .05.

**Figure S6.**
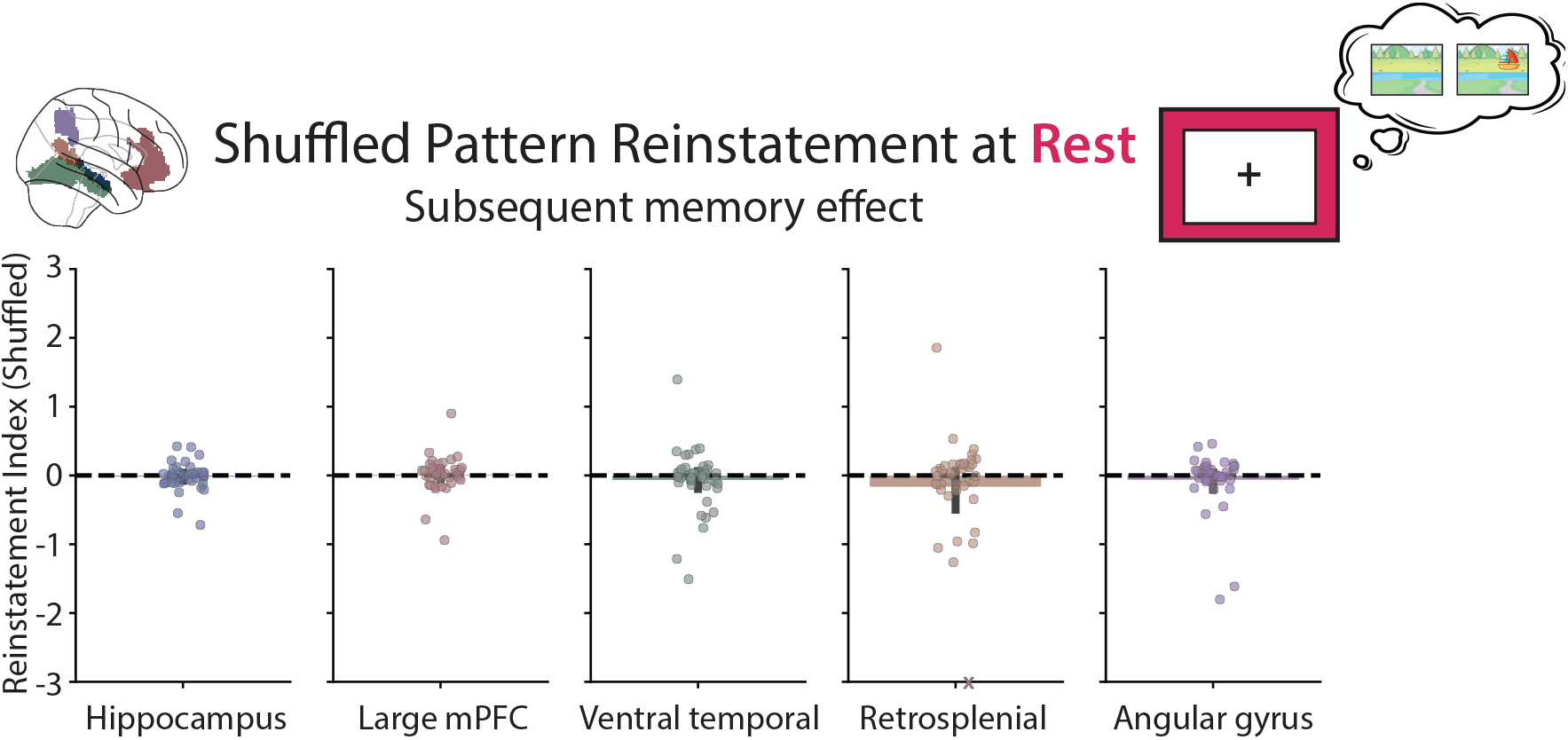
Spontaneous pattern reinstatement after random shuffling of labels. Remembered and forgotten labels were shuffled 1000 times within participant in anatomically defined ROIs. For each permutation, the reinstatement index was calculated as the difference in the summed correlation between the (shuffled) remembered trials minus the (shuffled) forgotten trials during post-encoding rest, after accounting for pre-encoding rest. These permuted reinstatement indices were then averaged within-participant before running group statistics. None of the ROIs showed a significant difference in the reinstatement index when the memory labels were shuffled (all *p*s > 0.266). Dots represent individual participant data and error bars reflect the 95% confidence interval from bootstrap resampling. Participants with values outside of the plotting range are marked with Xs on the negative or positive edge of the y-axis.

**Figure S7.**
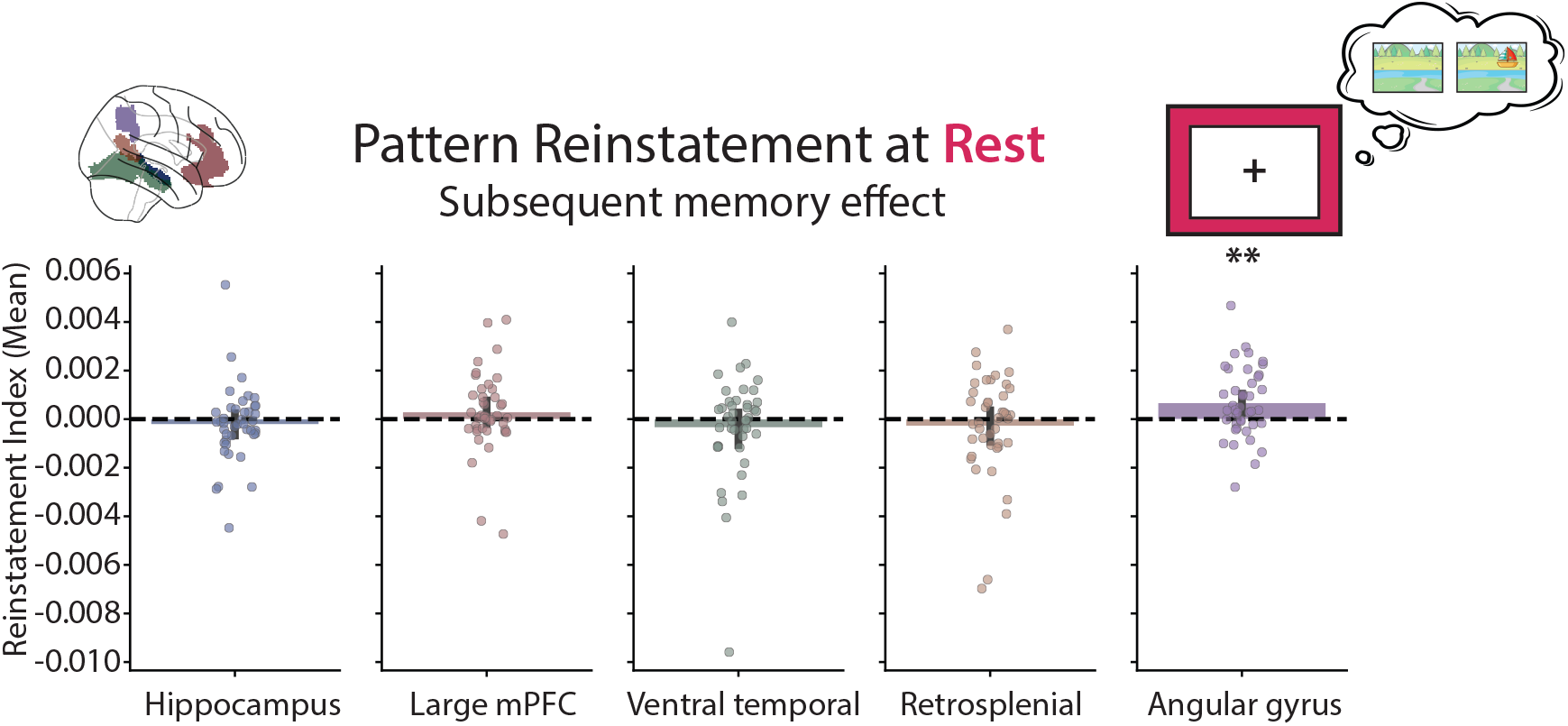
Spontaneous pattern reinstatement using the average correlation. Contrast of subsequently remembered versus subsequently forgotten trials during post-encoding rest in different anatomically defined ROIs when averaging correlations (rather than summing, as in the main analysis in Figure 3). Dots represent individual participant data and error bars reflect the 95% confidence interval from bootstrap resampling. ** *p* < .01.

**Figure S8.**
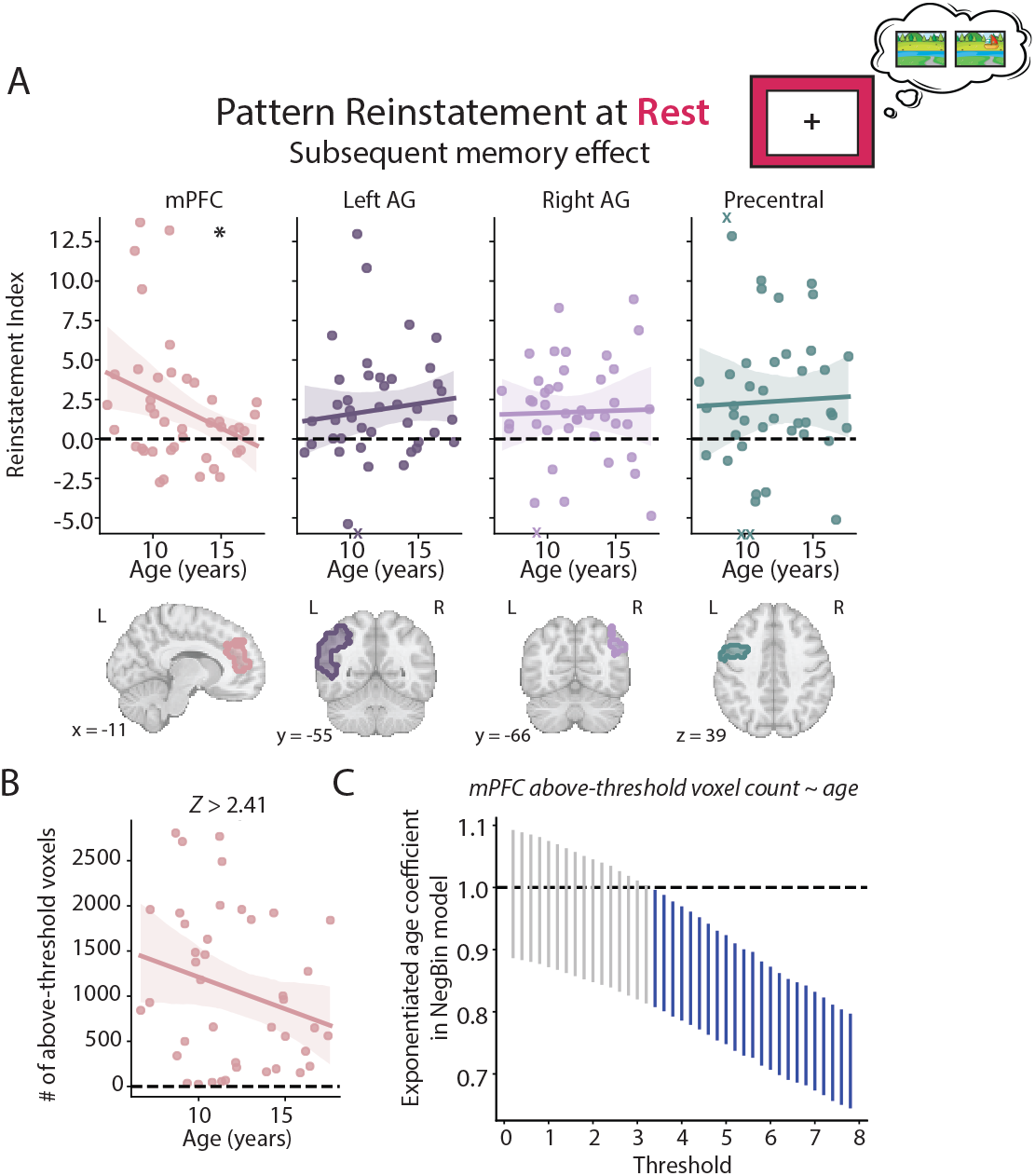
Effects of age on spontaneous pattern reinstatement. (**A**) Relationship between age and the contrast of subsequently remembered versus subsequently forgotten trials during post-encoding rest in regions significant in the whole brain (Figure 3**; Table S3**). There was a significant negative effect of age on pattern reinstatement in the mPFC (*r* = −.318, *p* = .040); no age effects were found in any of the other regions (all *p*s >.499). mPFC results are replotted from Figure 3B. Dots represent individual participant data and shaded error regions reflect the 95% confidence interval from bootstrap resampling. Participants with values outside of the plotting range are marked with Xs on the positive or negative edge of the y-axis. (**B**) Relationship between age and number of voxels above-threshold in the mPFC using the threshold from the whole-brain analysis (*z* > 2.41). In a negative binomial regression model predicting voxel count from age, there was a non-significant effect of age on voxel count (incidence rate ratio [IRR] = 0.93, CIs = [0.841 to 1.03], *p* = .174). (**C**) Confidence intervals for the exponentiated coefficient (IRR) for age in negative binomial regression models predicting above-threshold voxel counts at different thresholds. At higher thresholds, the IRR confidence intervals were below 1 (dark blue lines), indicating a negative effect of age on above-threshold voxel count. mPFC = medial prefrontal cortex, AG = angular gyrus. * *p* < .05.

**Figure S9.**
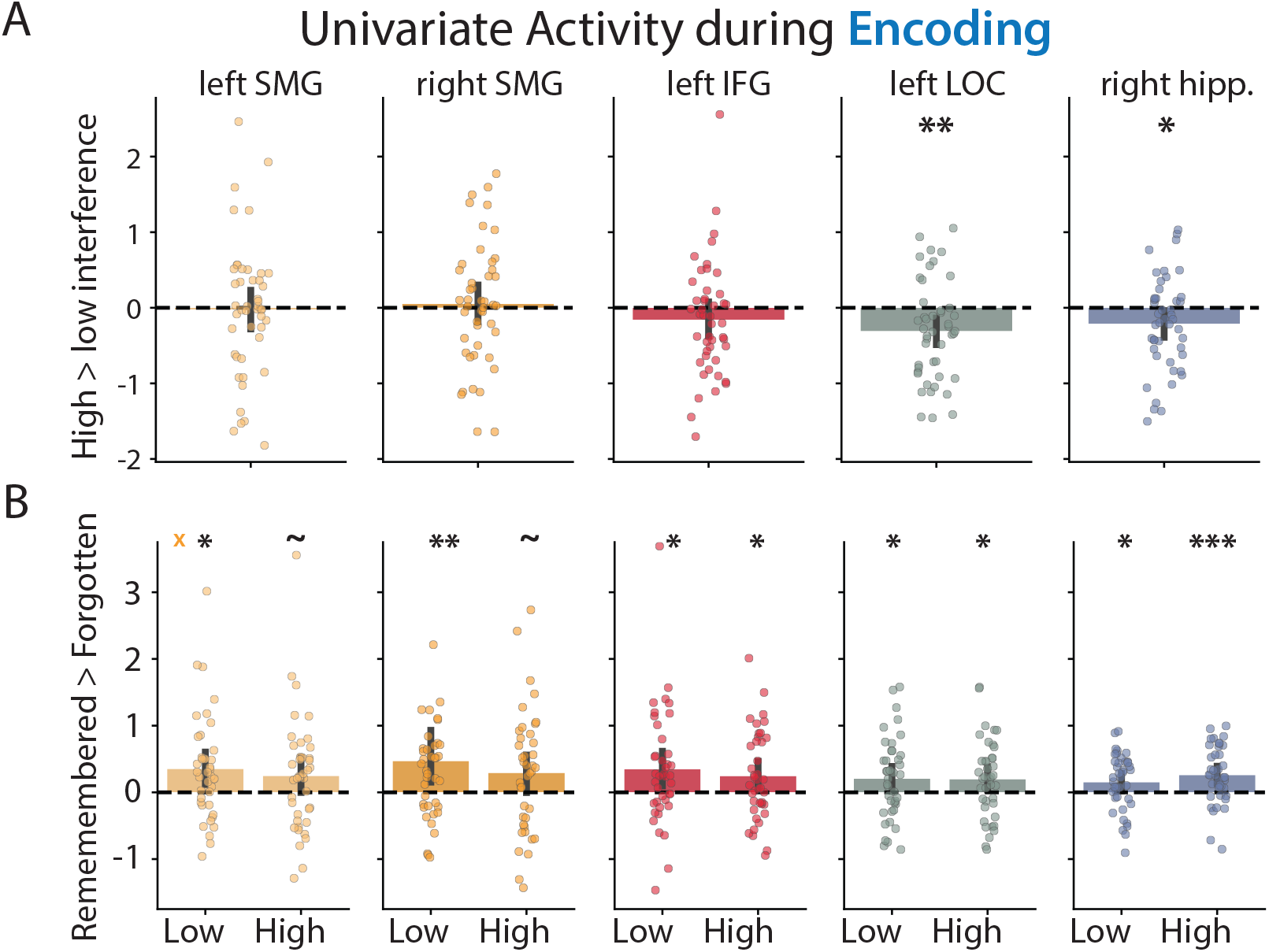
Effects of trial type on memory encoding activity. (**A**) Contrast of high versus low interference trials during memory encoding in regions that showed a significant subsequent memory effect in the whole brain. There was less activation in the left LOC (*M* = −0.31, *t*(45) = −3.21, *p* = .002, *d* = 0.48) and right hippocampus (*M* = −0.21, *t*(45) = −2.34, *p* = .024, *d* = −0.35) for high versus low interference trials; no other region showed an overall effect of trial type (left SMG: *M* = −0.03, *t*(45) = −0.19, *p* = .853, *d* = −0.03; right SMG: *M* = 0.05, *t*(45) = 0.43, *p* = .669, *d* = 0.06; left IFG: *M* = −0.16, *t*(45) = −1.42, *p* = .164, *d* = −0.21). (**B**) Contrast of remembered versus forgotten trials for low and high interference trials in a subset of participants with at least one trial in each condition. All regions showed a significant subsequent memory effect for low interference trials (left SMG: *M* = 0.35, *t*(39) = 2.85, *p* = .007, *d* = 0.46; right SMG: *M* = 0.47, *t*(39) = 2.29, *p* = .027, *d* = 0.37; left IFG: *M* = 0.35, *t*(39) = 2.51, *p* = .016, *d* = 0.40; left LOC: *M* = 0.21, *t*(39) = 2.12, *p* = .041, *d* = 0.34; right hippocampus: *M* = 0.15, *t*(39) = 2.27, *p* = .029, *d* = 0.36) and significant or trending subsequent memory effects for high interference trials (right SMG: *M* = 0.29, *t*(39) = 2.03, *p* = .050, *d* = 0.33; left SMG: *M* = 0.25, *t*(39) = 1.81, *p* = .078, *d* = 0.29; left IFG: *M* = 0.25, *t*(39) = 2.34, *p* = .025, *d* = 0.38; left LOC: *M* = 0.20, *t*(39) = 2.13, *p* = .039, *d* = 0.34; right hippocampus: *M* = 0.26, *t*(39) = 3.85, *p* < .001, *d* = 0.62). There was no difference in subsequent memory effects for low versus high interference trials across regions (all *p*s > .306). Dots represent individual participant data and error bars reflect the 95% confidence interval from bootstrap resampling. Participants with values outside of the plotting range are marked with Xs on the negative or positive edge of the y-axis. Hippocampal results are replotted from Figure 4. SMG = supramarginal gyrus, IFG = inferior frontal gyrus, LOC = lateral occipital cortex, hipp. = hippocampus. *** *p* < .001, ** *p* < .01, * *p* < .05, ∼ *p* < 0.10.

**Figure S10.**
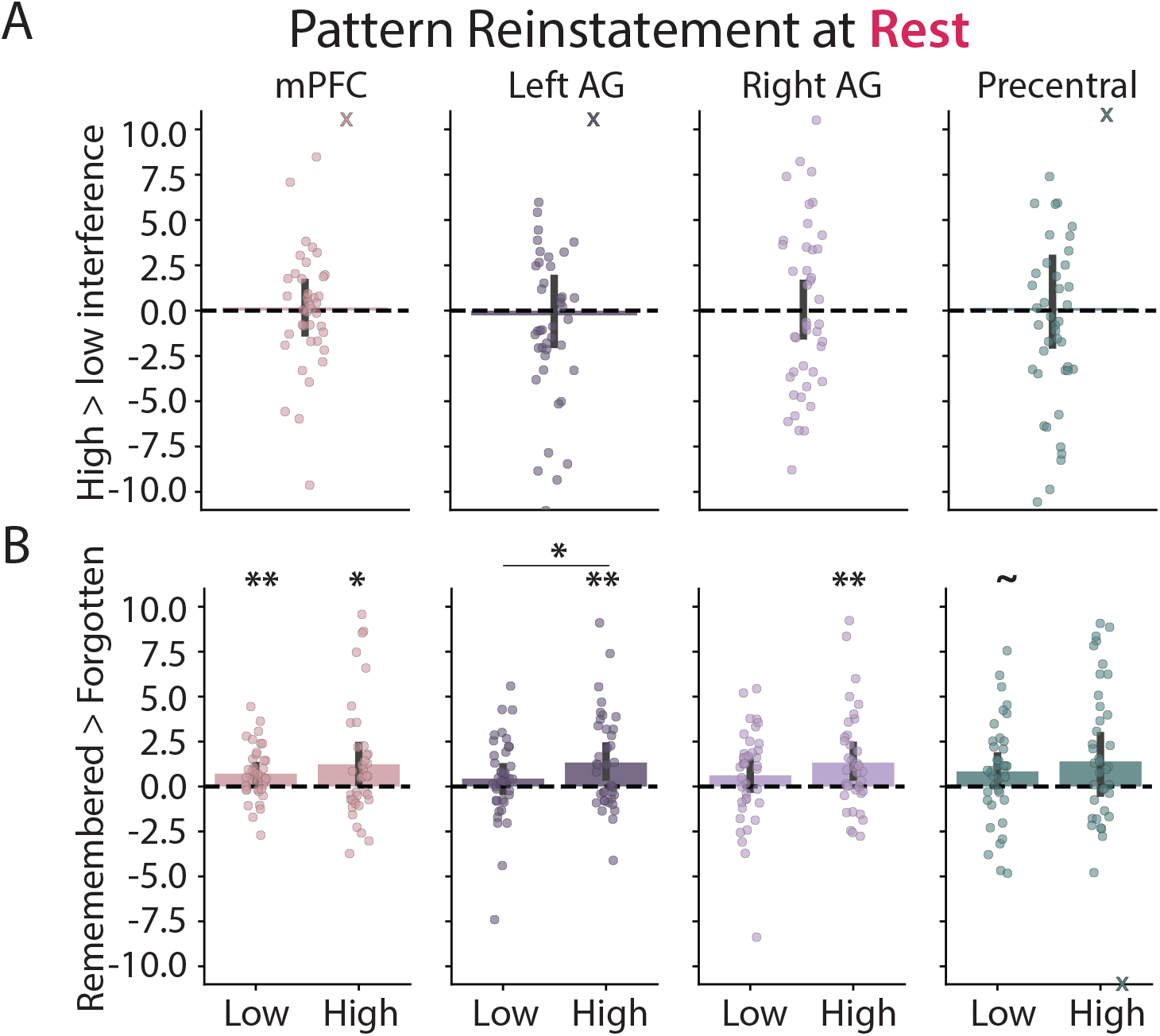
Effects of trial type on pattern reinstatement. (**A**) Contrast of pattern reinstatement for high versus low interference trials during post-encoding rest (after accounting for pre-encoding rest) in regions that showed significant pattern reinstatement in the whole brain. No region showed an overall effect of trial type (mPFC: *M* = 0.19, *t*(41) = 0.26, *p* = .795, *d* = 0.04; left AG: *M* = −0.27, *t*(41) = −0.28, *p* = .778, *d* = −0.04; right AG: *M* = 0.02, *t*(41) = 0.03, *p* = .976, *d* = 0.01; precentral: *M* = 0.16, *t*(41) = 0.12, *p* = .902, *d* = 0.02). (**B**) Contrast of pattern reinstatement for remembered versus forgotten trials for low and high interference trials in a subset of participants with at least one trial in each condition being tested. There was significantly greater pattern reinstatement of remembered versus forgotten low interference trials in the mPFC (*M* = 0.73, *t*(39) = 3.21, *p* = .003, *d* = 0.51), with marginal effects in the precentral cortex (*M* = 0.84, *t*(39) = 1.93, *p* = .062, *d* = 0.31) and no effects in the angular gyrus (left AG: *M* = 0.44, *t*(39) = 1.21, *p* = .234, *d* = 0.19; right AG: *M* = 0.64, *t*(39) = 1.57, *p* = .126, *d* = 0.25). For high interference trials, there was significantly greater pattern reinstatement of remembered versus forgotten trials in all regions except the precentral cortex (mPFC: *M* = 1.23, *t*(37) = 2.33, *p* = .025, *d* = 0.38; left AG: *M* = 1.35, *t*(37) = 3.08, *p* = .004, *d* = 0.51; right AG: *M* = 1.35, *t*(37) = 3.00, *p* = .005, *d* = 0.49; precentral: *M* = 1.41, *t*(37) = 1.69, *p* = .100, *d* = 0.28). There was significantly greater subsequent memory reinstatement for high versus low interference trials in the left AG (*t*(35) = 2.09, *p* = .044, *d* = 0.35); no other regions showed a difference in subsequent memory reinstatement by trial type (all *p*s > .211). Dots represent individual participant data and error bars reflect the 95% confidence interval from bootstrap resampling. Participants with values outside of the plotting range are marked with Xs on the negative or positive edge of the y-axis. mPFC results are replotted from Figure 4. mPFC = medial prefrontal gyrus, AG = angular gyrus. ** *p* < .01, * *p* < .05, ∼ *p* < 0.10.

**Figure S11.**
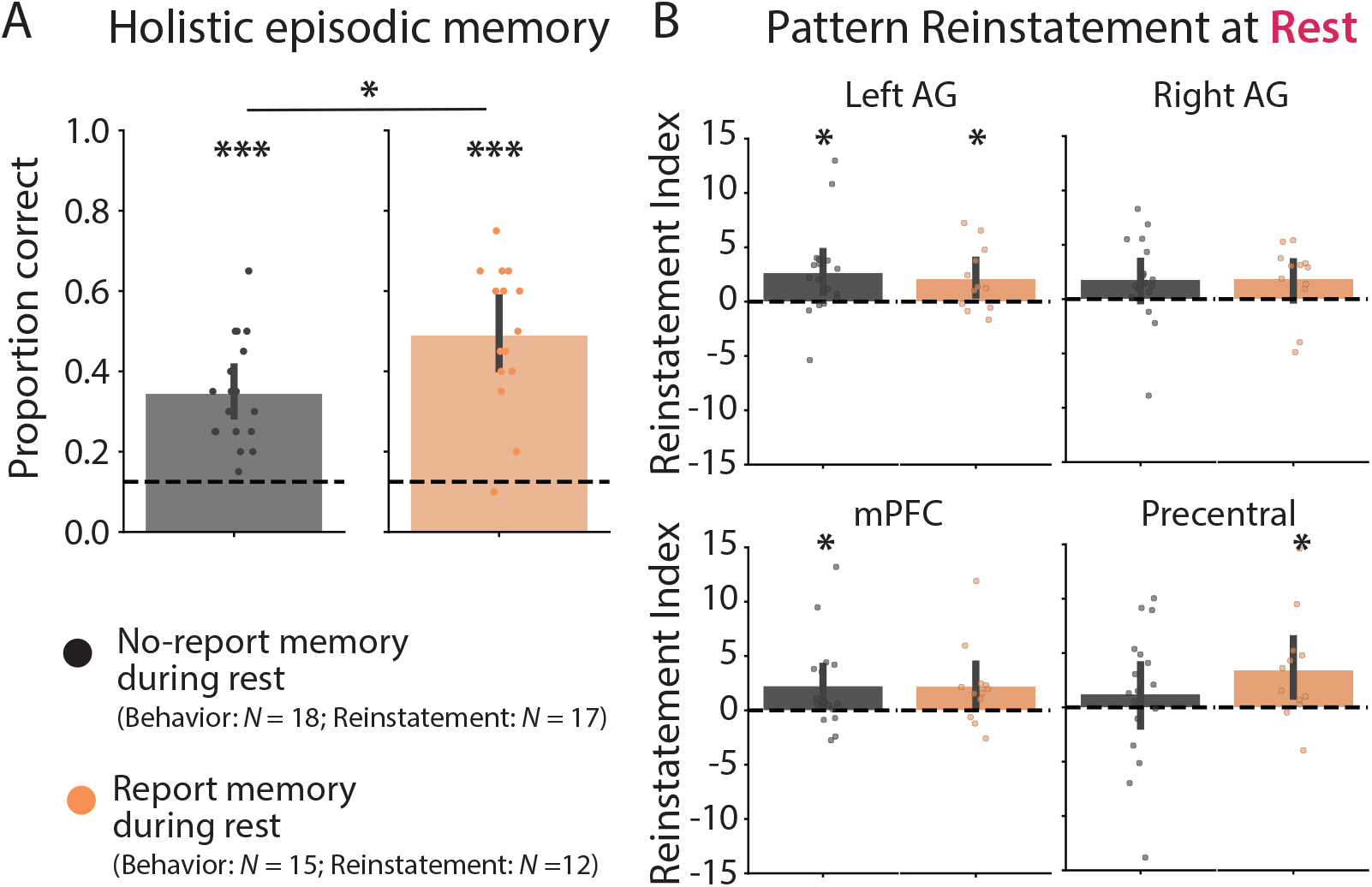
Effects of reported memory during rest on memory behavior and reinstatement. (**A**) Holistic episodic memory performance broken down by whether participants reported having thought about the memory task during post-encoding rest (orange) or not (dark grey) in participants who completed the debrief questionnaire (*N* = 33 out of 49 total for memory behavior analysis, *N* = 29 for reinstatement analysis). An ANCOVA with age and reporting group as predictors revealed a main effect of reporting group on memory performance (*F*(1, 30) = 7.09, *p* = .012, *η*_G_^2^ = .191). A follow-up *t*-test revealed better holistic episodic memory for participants who reported thinking about the memory task during the rest period (*t*(32) = 2.67, *p* = .012, d = 0.93). (**B**) Pattern reinstatement for remembered and forgotten trials in regions significant in the whole-brain searchlight analysis for participants who reported having thought about the memory task during post-encoding rest or not. No main effect of reporting group was found on memory reinstatement (all *p*s > .316). Dots represent individual participants and error bars represent 95% confidence errors from bootstrap resampling. mPFC = medial prefrontal gyrus, AG = angular gyrus. *** *p* < .001, * *p* < .05.

**Figure S12.**
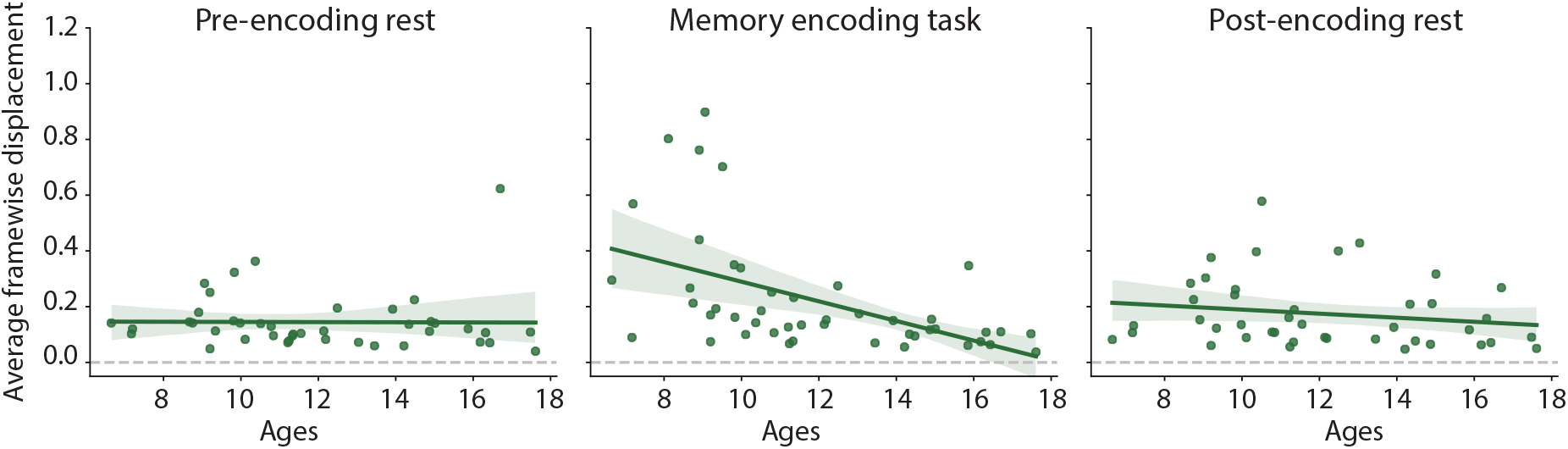
Relationship between motion and age. Average framewise displacement for the three fMRI runs (pre-encoding rest, memory encoding, and post-encoding rest) across all participants with usable data for that run (i.e., < 30% of timepoints with > 0.9 mm framewise displacement) and who were included in the analyses (N = 46 for memory encoding and N = 42 for pre-and post-encoding rest runs). For the memory encoding run only, average framewise displacement decreased over development (*r* = −.51, *p* < .001). Dots represent individual participants and shaded error regions represent 95% confidence errors from bootstrap resampling.

